# From SCUBA to spectra: Broadly applicable methods for coral metabolomics research

**DOI:** 10.64898/2026.01.19.700417

**Authors:** Sabrina L. Rosset, Thomas J.B. Cline, Kiran-Kumar Shivaiah, Immy Ashley, Khalil Smith, Giada Tortorelli, Rapid Resilient Reefs Consortium, Lauren K. Walling, John E. Parkinson, Mark J.A. Vermeij, Mehdi Bouhaddou, Crawford Drury, Ty N.F. Roach, Robert A. Quinn

**Author notes:** Detailed author list in supplementary file (Rapid Resilient Reefs Consortium).

## Abstract

Corals represent a complex assemblage consisting of a host cnidarian, symbiotic dinoflagellate microalgae, and associated microbiomes and viromes, collectively called the coral holobiont. Corals are foundational to tropical reefs, yet their global decline due to climate change and other stressors creates an uncertain future for this valuable ecosystem. Metabolomics is a powerful means to unravel biochemical interactions within the holobiont that underpin coral resilience and adaptation. However, the remote nature of reefs and the analytical demands of this technique often limit its application. Untargeted metabolomics presents analytical challenges that are amplified in complex samples like corals, such as identifying the biological source of metabolites. Here, we evaluate how different sample fixation methods and time delays before storage—unavoidable in field contexts—affect coral metabolome profiles. We further present a framework for mapping metabolites in holobiont samples to their coral host and algal symbiont origins and introduce a spectral library to improve and automate annotation of coral lipids. Additionally, we demonstrate how single samples can be used concurrently for metabolomics, DNA amplification, and proteomics. Together, our study provides a streamlined, field-adaptable workflow for coral metabolomics that enables larger-scale studies and broader adoption of metabolomics in coral reef research and conservation.

## Introduction

Coral reefs face escalating threats from anthropogenic pressures, particularly ocean warming, which threatens the long-term functioning of this ecosystem [1, 2]. The coral–algal mutualism is a nutritional symbiosis in which photosynthetically derived sugars and lipids produced by the algae are transferred to the host to support calcification and growth [3]. This metabolic interaction forms the energetic foundation of coral reefs but also represents a point of vulnerability. Stressors such as excess heat and light can destabilize the association and cause symbiont loss in the form of coral bleaching, which can lead to colony mortality and reef degradation [4]. Gaining a better understanding of the metabolic interactions that sustain the symbiosis, especially under stress, is critical [5, 6]. Rapid advancements in metabolomics instrumentation and bioinformatic tools [7, 8] provide an exciting avenue for accelerated discovery and the potential to support science-based coral reef conservation and restoration strategies [9].

Metabolomics is an analytical technique to profile, characterize and quantify small molecules in biological samples. Comprehensive analysis of the metabolome – the complete set of small molecules (typically < 1,500 Daltons) – can provide extensive insight into the physiological and metabolic state of an organism and reveal biochemical responses to microbial or environmental interactions [10]. Because of the complex nature and vast biochemical diversity of metabolomes, no single method can capture it entirely. This complexity favors untargeted metabolomics, which aims to broadly profile relative abundances of compounds for hypothesis-generation or biomarker discovery using high-resolution mass spectrometry [11]. Diverse sample preparation procedures, chromatographic conditions and analytical platforms have been developed to capture various components of the metabolome, including targeted methods to detect and quantify specific known compounds of interest. There is an inherent tradeoff between methods that most broadly or accurately capture metabolites *versus* the ease and high-throughput nature of their application; considerations that weigh heavily in systems where sample collection itself is challenging, such as coral reefs.

Multiple studies have conducted metabolomics analysis of reef-building Scleractinian corals collected from field and laboratory settings. This body of work has characterized the biochemical diversity of corals and their algal symbionts [12–18], and revealed molecular responses or adaptations to symbiosis [19], heat stress [20–27] and disease [28]. For example, symbiont membrane composition [21, 29, 30] and coral lipid reserves [31, 32] have been highlighted as potential mechanisms contributing to coral thermal resilience. However, the metabolic basis of (1) coral–symbiont–microbe interactions, (2) holobiont thermal tolerance, and (3) holobiont adaptive capacity remains incompletely understood and warrants further investigation.

Coral metabolomes are shaped by complex interactions between the coral host, its algal symbionts (dinoflagellates in the family Symbiodiniaceae), other microbes, and the environment. Holobiont biochemistry is profoundly affected by the dominant microalgal partner and hypothesized to influence thermal tolerance [30, 33, 34]. A unique challenge in metabolomics, absent in sequencing-based techniques where organismal identity is inherent in the nucleic acid or protein sequence data, is identifying the origin of metabolites in complex communities and holobionts [35, 36]. In corals, disentangling host- and symbiont-derived metabolites typically requires laborious sample preparation and separation steps that limit throughput [37].

Metabolomics studies of corals have used diverse analytical strategies tailored to specific research objectives, including gas-chromatography-mass spectrometry (GC-MS) to analyze primary metabolites and fatty acids [25, 38–41], or targeted liquid-chromatography–tandem mass spectrometry (LC-MS/MS) to profile specific lipid classes [21, 42, 43]. Increasingly, untargeted LC-MS/MS is being employed to report both the annotated and unannotated chemistry of corals [16, 20, 22, 30]. Despite growing research interest in this area, compound annotation remains a challenge, and the underrepresentation of coral metabolites in public spectral libraries requires substantial manual curation and expertise in biochemistry to achieve accurate annotations [24, 44].

Here we present practical guidelines for sampling corals for downstream metabolomic analysis. Using diverse coral samples collected from the field and aquaria, we evaluate the effects of different sample fixation and handling methods, develop a method for mapping metabolites in holobiont samples to their host and symbiont compartments, establish a spectral library of coral lipids, and demonstrate the compatibility of metabolomics extracts with DNA amplification and proteomics. Together, this work aims to provide a streamlined, field-adaptable framework for coral metabolomics analysis that facilitates broader adoption of this technique and can be adapted for large-scale studies across reef systems worldwide.

## Materials and Methods

For this study and method development, we included three sample sets: 1) corals collected from reefs in Curaçao at the Caribbean Research and Management of Biodiversity Institute (CARMABI), 2) corals held in flow through aquaria at the Hawaiʻi Institute of Marine Biology (HIMB) and 3) *Galaxea fascicularis* held in closed aquaria at Michigan State University.

### Field collection methods for coral biopsies

On reefs in Curaçao and Kāneʻohe Bay, Hawaiʻi, we developed methods for consistent sample collection based on coral colony morphology and skeleton density using either bone clippers (*e.g.*, BRS SPS Coral Bone Cutter) or a steel dermal curette (*e.g.*, 5 mm, round dermal curette; Figures 1 and S1). Mounding and encrusting species with non-meandroid polyps (*e.g., Orbicella faveolata, Montastrea annularis*) were sampled directly with the curette. Branching species (*e.g.*, *Acropora cervicornis, Pocillopora acuta)* and laminar/foliose species (*e.g., Pavona varians*) and mounding/encrusting species with meandroid polyps (*e.g., Diploria labrinthiformis)* were sampled with clippers before subsampling with the curette. Plating species were sampled by clipping a pie-shaped piece from the edge of the plate, while meandroid species were sampled by clipping septae/ridges between polyps.

**Figure 1.**
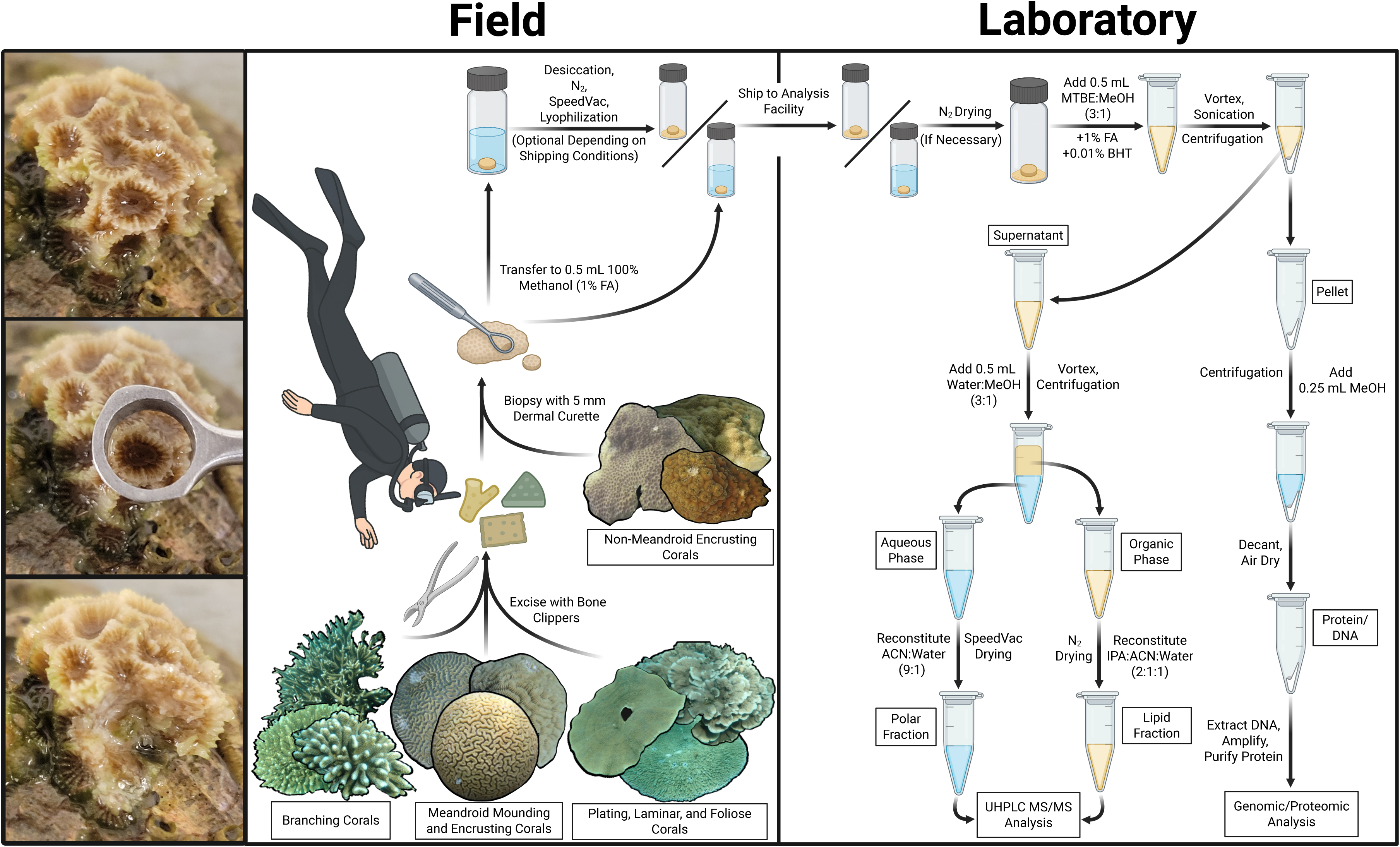
Workflow for field-based coral sample collection and fixation and laboratory-based extraction for metabolome analysis with dual genomic/proteomic profiling. In the field, coral biopsies are collected using either a 5 mm dermal curette or bone clippers, depending on colony morphology. An example of sample collection with a dermal curette is shown for *Astrangia poculata,* where the curette is centered over a single polyp (left-hand panels). Coral biopsies are optimally fixed by transfer into a glass vial containing ice-cold, acidified methanol (1% formic acid). Samples can optionally be desiccated for more stable shipment to laboratories if cold shipment or shipment of methanol is unavailable. Once in a laboratory, the methanol is evaporated unless desiccated already, and samples are then extracted in MTBE:methanol (3:1) by thorough vortexing and sonication. The pellet obtained after centrifugation can be used for genomic and/or proteomic analysis. The supernatant is further extracted by adding an equal volume of water:methanol (3:1) for liquid:liquid phase separation into aqueous and organic phases that are collected, dried and reconstituted in appropriate solvents for the analysis of polar metabolites and mid-to-non-polar metabolites (mostly lipids), respectively, by untargeted ultra-high-pressure-liquid chromatography – tandem mass spectrometry (UHPLC-MS/MS) using a high resolution mass spectrometer. Figure created in BioRender. Cline, T. (2026) https://BioRender.com/4jr2kld.

When diving, samples were stored in labeled ziplock bags containing seawater from the collection site. Upon surfacing, they were placed in a >20-gallon cooler filled with ambient seawater, where they remained until extraction. If necessary, coral fragments were subsampled by placing larger fragments on a methanol-cleaned, flat surface and positioning the curette over the center of a polyp before pressing into the tissue and skeleton (Figure S1). All biopsies were collected deep enough to retrieve tissue and skeleton regardless of colony morphology. Using forceps, biopsies were briefly blotted on tissue paper to remove excess sea water and then placed directly into solvent (as described below – Figure S1). All samples were collected under the collecting permits of the Curaçaoan Government provided to CARMABI (2022/021467) and SAP 2022-22 to HIMB.

### Coral sample treatment, storage, extraction, and mass spectrometry

#### Sample collection and treatment

To test the effects of different metabolome quenching and sample handling methods on the coral holobiont metabolomic profile, we collected replicate biopsies of corals maintained in flow-through tanks at HIMB and subjected the samples to different treatments. Biopsies were collected from four species of corals to explore any species-specific differences in treatment outcomes. Biopsies of *Montipora capitata*, *Montipora patula* and *Porites compressa* were collected via the dermal curette technique while biopsies of *Pocillopora meandrina* were collected with bone clippers. Samples were collected from five colonies per species (n = 5 replicates per treatment). Treatments were as follows: (1) biopsies were transferred to cryovials and flash frozen in liquid nitrogen; (2) biopsies were transferred to small amber glass vials containing 500 μL of 100% ethanol, (3) 500 μL of 100% HPLC-grade methanol or (4) 500 μL of methanol with added 1% formic acid; and finally (5) biopsies were transferred to cryovials and heated to 95°C for 5 min in a heat block before being stored on ice. The latter heating method was tested due to its common application in proteomics whereby enzymatic activity is rapidly quenched by protein denaturation.

During collection, all vials were pre-chilled and stored in a Styrofoam box with ice; they were transferred to a -80°C freezer 1 h after collection. Additionally, for a subset of samples quenched in either 100% methanol or acidified methanol (1% formic acid), we tested the effect of temporary storage duration (the time delay from sample collection until safe storage in a - 80°C freezer). Biopsies were stored on ice (∼4°C) for 1 h, 4 h, 8 h or 24 h until transfer to the freezer. All samples were shipped to Michigan State University on dry ice (48 h shipment) for mass spectrometry analysis. Finally, for one set of samples fixed in 100% methanol (1 h until transfer to freezer), we tested the effect of storing at room temperature for 48 h after drying the biopsy completely under a stream of N_2_ gas to simulate the option of shipping samples to a lab without access to dry ice or other forms of temperature-controlled shipping.

#### Metabolite extraction

The extraction and data acquisition methods chosen depend on the specific research question and molecules or pathways of interest. Here we used a method that captures a broad spectrum of the metabolome by conducting a two-phase extraction using methanol and methyl-tert-butyl-ether (MTBE), adapted from Salem et al. [45]. The resulting aqueous phase is analyzed using hydrophilic interaction liquid chromatography to profile small polar metabolites, such as amino acids, peptides or methylated amines, while the organic phase is analyzed using C18 reverse phase chromatography to profile lipids (primarily) and other compounds ranging from mid-to-high hydrophobicity (see detailed protocol in Supplementary Materials).

To assess the effect of sample treatment on the coral metabolomic profile, we analyzed the organic “lipid” fractions only. Briefly, each sample was first dried under a stream of N_2_ gas to remove any inconsistency in solvent volumes due to evaporation before being extracted in 500 μL of ice-cold MTBE:methanol (3:1, v:v) containing 1% formic acid and 0.01% butylated hydroxytoluene (BHT) by vortexing thoroughly, sonicating in a water bath sonicator with ice for 15 min, and then vortexing again. Formic acid and BHT served to prevent metabolite degradation during the extraction process by inhibiting lipase activity and preventing lipid oxidation, respectively. The extract was then transferred into a 1.7 mL microcentrifuge tube (Axygen™ MCT-175-L-C), leaving the skeletal fragment behind, and centrifuged at 15,000 x g for 10 min at 4°C to pellet protein, DNA and cellular debris. The supernatant was transferred to a new 1.7 mL microcentrifuge tube before adding 500 μL of ice-cold water:methanol (3:1, v:v). The sample was vortexed thoroughly and then incubated at -20°C for 30 min to allow for phase separation before centrifuging at 10,000 x g for 10 min at 4 °C. The upper organic phase was carefully transferred into a new amber glass vial (Restek™ 21143), and the solvent was evaporated under a stream of N_2_ gas. The dried sample was then reconstituted in 500 μL of 2-propanol:acetonitrile:water (2:1:1, v:v:v). A quality control (QC) sample was prepared by pooling 20 μL of each sample into a new vial. All samples were stored at -80°C until analysis. Samples were kept on ice throughout the extraction protocol, apart from solvent evaporation steps, which were carried out at room temperature. Extraction blanks were prepared alongside the samples during the extraction protocol (n = 5). All solvents used were HPLC-grade.

#### Mass spectrometry data acquisition

Mass spectrometry analysis was conducted using a Vanquish ultra-high-performance liquid chromatography (UHPLC) system coupled to a Q Exactive mass spectrometer (Thermo Scientific). We note that other high resolution mass spectrometers are also suitable for data acquisition, but data-dependent MS/MS acquisition is essential for downstream data analysis (see below). The extracts were separated using a 20 min chromatography method at a flow rate of 0.4 mL/min, using a reverse phase Acquity UPLC BEH C18 column (130 Å, 1.7μm, 2.1 x 100 mm, Waters) fit with a VanGuard pre-column (Acquity UPLC BEH C18, 130 Å, 1.7μm, 2.1 x 5 mm, Waters) as the stationary phase and the following mobile phases: solvent A (60:40 acetonitrile:water with 0.1% formic acid and 10 mM ammonium formate) and solvent B (90:10 2-propanol:acetonitrile with 0.1% formic acid and 10 mM ammonium formate). Data was acquired in positive ionization mode using data-dependent acquisition. Detailed chromatographic and mass spectrometry parameters are provided in the Supplementary Materials. An 8-point standard curve of the QC sample was acquired in triplicate during the run (0.1, 0.2, 0.4, 0.6, 0.8, 1.0, 1.2, and 1.6-times QC concentration).

### Metabolomics data processing and analysis

#### Data processing

Metabolite feature detection and alignment were performed using MZmine software (v 4.3; Table S5) [46]. Non-biological features were removed from the dataset by discarding features with 3x higher mean signal intensity in extraction blanks compared to samples and QCs.

Sample quality was assessed by plotting richness (*i.e.,* total number of detected features) *versus* maximal signal intensity of all samples and QCs to identify potential variability in metabolome acquisition depth across the sample set (as in Figure 4c), as well as by data visualization in PCA ordination and by performing an outlier analysis using Mahalanobis distance and the Isolation Forest (IF) method in the ‘*solitude’* package [47]. Samples with low richness and signal intensity (∼ <20% of QC mean) were removed from the dataset, as were those identified as outliers using the IF method. Features that were detected in <5 samples per colony (out of 10 samples total – 1 per treatment) were removed as an additional step to reduce noise. The data were then normalized using the maximal density fold change (MDFC) post-acquisition normalization method in the ‘*MAFFIN’* package [48]. Note that project needs may favor probabilistic quotient normalization (PQN) normalization. The data were then filtered using high quality feature selection in the ‘*MAFFIN’* package using a relative standard deviation threshold in QC samples of 0.4 and a Pearson correlation to serial QC samples threshold of 0.8 [49]. Compound annotation was performed as described below.

#### Data analysis

All data analysis was performed in R (v 4.5.0). To assess how coral holobiont metabolomic profiles were influenced by different metabolome quenching and sample handling methods (liquid nitrogen, ethanol, methanol, acidified methanol, boiled, dessicated-48 h RT), we applied permutational multivariate analysis of variance (PERMANOVA) and partial least squares discriminate analyses (PLS-DA) based on Bray-Curtis dissimilarities for each of the four coral species analyzed. We identified metabolite classes that were most sensitive to sample treatment using Wilcoxon Rank Sum tests to extract those classes that significantly clustered towards high PLS-DA loading scores in components 1 and 2 (*p* < 0.05, FDR-adjusted). Significant classes were visualized in heatmaps as the z-scaled summed abundance values of all features within the respective class. To assess the effect of prolonged storage of methanol-fixed samples on ice prior to transfer to a freezer, we calculated Spearman correlation coefficients of all metabolite features to time (1h, 4 h, 8 h, 24 h) irrespective of coral species; again, we identified significantly correlated metabolite classes using Wilcoxon Rank Sum tests (*p* < 0.1, FDR-adjusted). The time-dependent rate of sample degradation was compared between samples quenched in acidified *versus* non-acidified methanol. The significance of the main effects of time and acidification was determined for each species using two-way analysis of variance (two-way ANOVA). Additionally, we evaluated the impact of sample fixation method on metabolite feature detection and abundance by assessing metabolome diversity metrics (richness and evenness) using the ‘*vegan’* package in R [50], and by calculating the coefficient of variance (%CV) for each high-quality metabolite feature across biological replicates within each treatment group (CV = 100 × SD/mean).

### MS/MS library curation and holobiont metabolite mapping

#### Compound annotation and curation of a spectral library of coral lipids

To better facilitate annotation of coral metabolites with our untargeted LC-MS/MS method, we used a broadly representative collection of Hawaiian coral species as well as a wide phylogenetic spread of algal symbionts (isolated cultures) to create a library of MS/MS spectra for public use. We performed extensive manual annotation of a dataset consisting of eight species sampled from across Kāneʻohe Bay, Hawaiʻi, with 15 colonies per species of *Montipora capitata*, *Montipora patula*, *Porites compressa*, *Porites lobata*, *Pavona varians*, *Leptastrea purpurea, Pocillopora acuta and Pocillopora meandrina* (SAP 2022-22 to HIMB). Holobiont biopsies from symbiotic adult colonies were fixed in 100% methanol. Additionally, this dataset was processed along with pooled samples made up of extracts derived from 115 Symbiodiniaceae cultures representing multiple species from five genera: *Symbiodinium*, *Breviolum, Cladocopium, Durusdinium* and *Effrenium.* Cultures were grown in F/2 liquid medium in an incubator maintained at 26°C with a 12:12 light:dark photoperiod and light intensity of ∼50 µmol quanta/m^2^/s as described by [51]. All samples were extracted and analyzed as described above. Together, this dataset represents a broad spectrum of coral and dinoflagellate biochemical diversity with which to create a spectral library.

Metabolite features were first annotated by matching their precursor masses (MS^1^) and fragmentation spectra (MS^2^) to libraries that are publicly available on the GNPS platform using a precursor cosine score cutoff of > 0.85 and a minimum of 4 matched fragments [52]. Additionally, features were assigned to compound classes based on *in silico* predictions in the CANOPUS module of SIRIUS [53, 54]. We then curated annotations of diverse lipid classes by manually annotating the spectra. To do this, we built custom functions in R to extract metabolite features from the spectral file of the dataset (.mgf file from mzMine output) containing specific product (P) or neutral loss (NL) masses that are diagnostic of specific membrane lipid classes (mz error < 0.008; Table 1) [55]. We then searched extracted precursor masses against the LipidMaps database (mz error < 0.005) [56] and against a custom database of predicted masses based on different fatty acid compositions, molecular adducts ([M+H]^+^, [M+Na]^+^ or [M+NH_4_]^+^) and oxidation states (mono-, di-, or tri-hydroxylated) for various lipid classes. We verified that the retention time of a feature matched the expected range for a given lipid class, for example, a shift to earlier retention times for oxidized *versus* non-oxidized lipids [57], and manually annotated exact fatty acid constituents where possible based on the MS^2^ spectra. By this approach, we identified and annotated multiple classes of phospholipids (PC, PE, PS, PI), phosphoshingolipids (SM, CerPE, CerPI, CAEP), glycerolipids (MGDG, DGDG, HexCer), sulfolipids (SQDG) and betaine lipids (DGCC, DGTS/A; see Table 1 for abbreviation definitions). By the same approach, we also annotated glycerolipids (MAG, DAG, TAG, MADAG) by identifying features with specific NL masses corresponding to fatty acids (Figure S4) [58]. Our annotation process was guided by available coral and Symbiodiniaceae lipidome datasets [17, 21, 24, 44, 59–63]. The resulting annotations were compiled into a spectral library and added to the publicly available database on the GNPS platform.

**Table 1.**
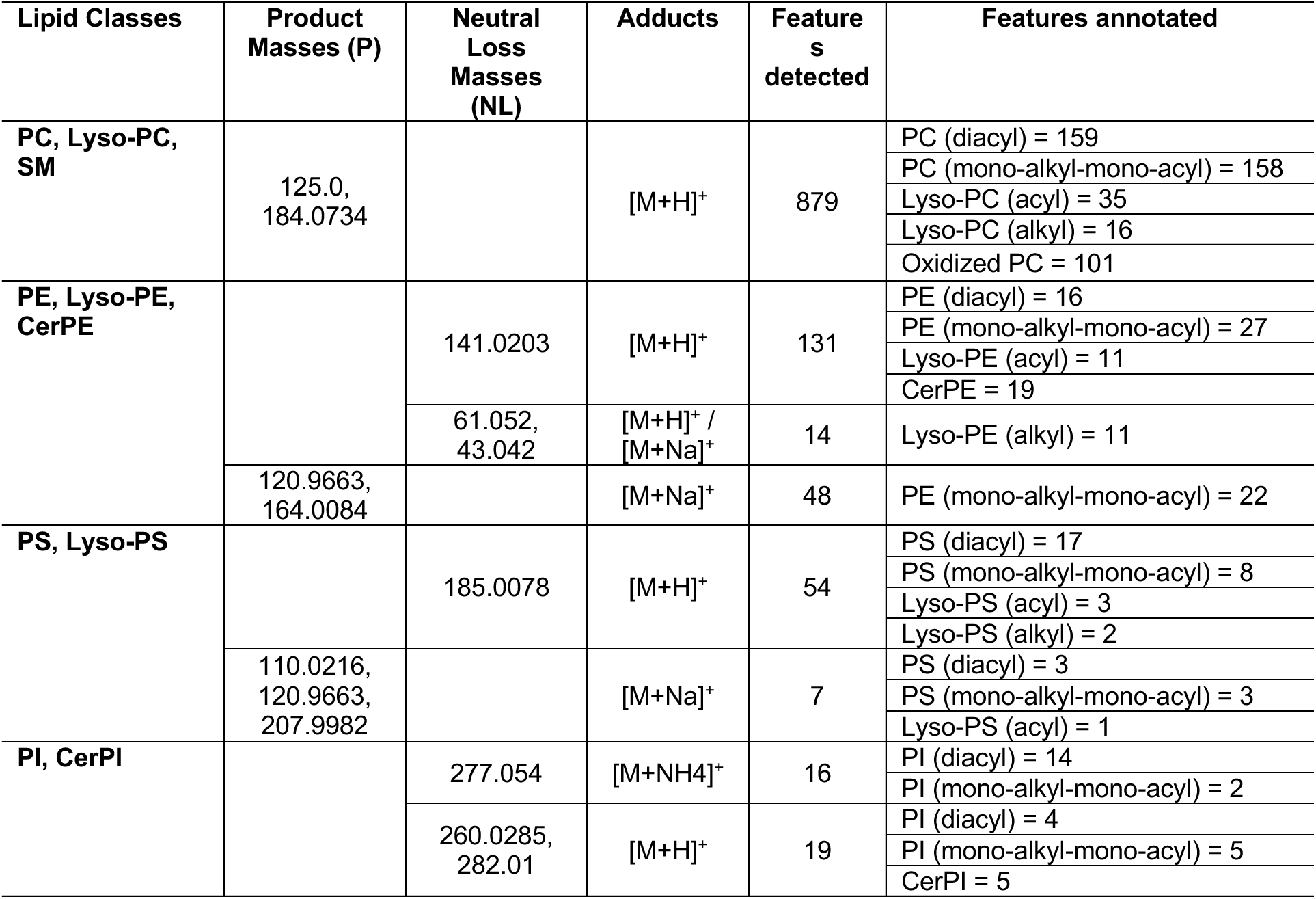
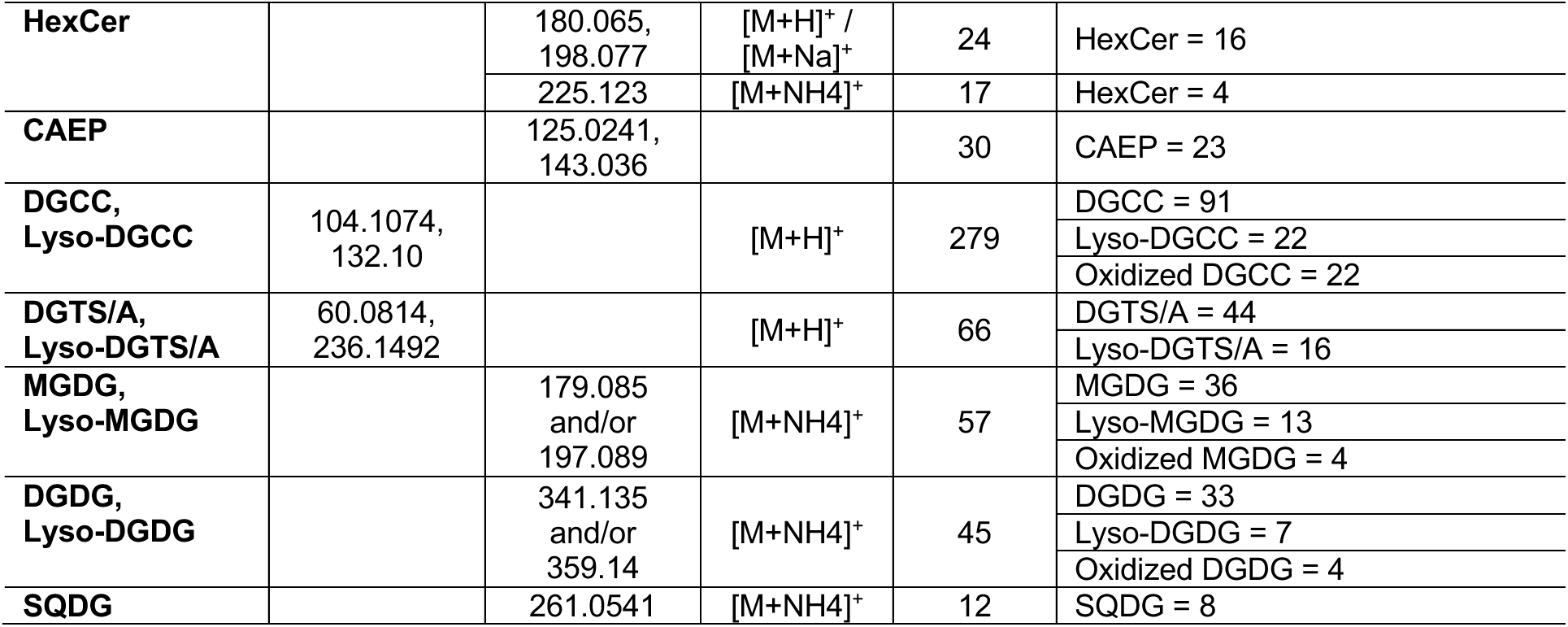
Characteristic product (P) and neutral loss (NL) masses used for lipid class identification, resulting numbers of detected features and those annotated as specific sub-classes. PC: phosphocholines, SM: sphingomyelins, PE: phosphoethanolamines, CerPE: ceramide phosphoethanolamines, PS: phosphoserines, PI: phosphoinositols, CerPI: ceramide phosphoinositols, HexCer: hexosylceramides, CAEP: ceramideaminoethylphosphonates, DGCC: diacylglyceryl carboxyhydroxymethylcholines, DGTS/A: diacylglyceryl trimethyl-homoserine/alanine, MGDG: monogalactosyldiacylglycerols, DGDG: digalactosyldiacylglycerols, SQDG: sulfoquinovosyldiacylglycerols

#### Metabolite mapping to coral host or algal symbiont origin

To determine the source of specific biochemicals (coral host or algal symbiont compartments), we processed (1) healthy, pigmented corals (multi-species dataset employed in spectral library generation), (2) experimentally bleached corals, (3) symbiont cultures, and (4) purified symbiont pellets derived from freshly isolated host tissues (see Supplemental Materials for detailed methods). Representative bleached tissues from each coral species were obtained by sampling corals following experimentally-induced thermal bleaching in temperature-controlled aquaria tanks at HIMB. Fully bleached coral samples were identified by the complete absence of three abundant algal lipids: DGCC(16:0/22:6), MGDG(18:3/18:5), and DGDG(18:4/20:5).

We assigned metabolite features to either coral host, algal symbiont, both host and symbiont, or unknown origin categories by assessing feature presence/absence and relative abundance across these samples. For each metabolite, relative abundances for each sample type were calculated and quasi-binomial generalized linear models were fit to compare abundances in symbiont culture *versus* bleached coral, as well as isolated symbiont pellet *versus* healthy coral. Effect sizes and significance were extracted alongside the number of samples with zero detection per group. Metabolites were classified as “coral host-derived” if present in bleached corals but absent in symbiont cultures and depleted in symbiont pellets; “algal symbiont-derived” if they were abundant in cultures and pellets but absent in bleached hosts; “both” if detected across all groups without strong enrichment. Metabolites were designated as “unknown” origin when their presence–absence patterns or relative abundances were inconsistent across reference sample types, when signal intensities fell below detection limits in multiple groups, or when they did not meet the predefined criteria for host, symbiont, or shared origin. A linear model was fit to test the correlation between the two enrichment patterns across all metabolite features (culture *versus* bleached host and isolated symbiont pellet *versus* healthy coral).

Origin assignments were further refined by applying network propagation of origin assignments to an MS/MS molecular network generated using the feature-based molecular networking workflow in GNPS [64]. The advanced diffusion algorithm was applied using the RCy3 package in R for two rounds of heat diffusion using time set to 1 [65]. Following each diffusion round, features with a diffusion score > 0.5 for either host or symbiont origin were re-assigned to their respective origin.

Finally, we tested if the resulting organismal origin assignments could be applied to a new dataset by adding the coral/algal designations of all features with their corresponding confidence score to our custom spectral library. We then used this library to provide taxonomic annotations to a dataset of healthy and menthol-bleached *Galaxea fascicularis* samples (see Supplementary Materials) using a cosine score cutoff of 0.65 and a minimum of 3 matched peaks in GNPS. We assessed the coverage and accuracy of origin assignments and compared them to results obtained using the same origin mapping framework described above using the Symbiodiniaceae culture samples along with extracts of symbionts isolated from the same *G. fascicularis* colonies and the menthol-bleached host samples.

### Multi-omics data integration – ITS2 amplification and proteomics

In an effort to obtain maximal information from a single coral biopsy, we tested the utility of our metabolite extraction method to allow for downstream nucleotide or peptide analysis of algal internal transcribed spacer 2 (ITS2) sequences and coral proteins. The pellet recovered following the first centrifugation step of the metabolite extraction (Figure 1) contains these biopolymers for downstream analysis.

#### DNA extraction and ITS2 amplification

We compared the efficiencies of symbiont ITS2 amplification between DNA extracted from precipitated DNA/protein pellets recovered from the metabolite extraction and DNA extracted from whole coral tissue biopsies in a typical workflow. For this purpose, we sampled paired biopsies of healthy coral tissue from *M. capitata*, *P. compressa,* and *P. acuta* fragments held in flow-through aquaria at HIMB (n = 10 colonies per species). One set of biopsies was flash frozen in liquid nitrogen for DNA extraction only, while the second set was fixed in 100% cold methanol and subjected to the metabolite extraction protocol described above. The precipitated pellets recovered from the metabolite extraction were first washed in 500 μL of cold 100% methanol, without disrupting the pellet, by centrifugation at 15,000 x g for 10 min at 4 °C and then discarding the supernatant to remove any residual MTBE. Washed pellets were air dried for 15 min to evaporate any remaining methanol and then re-suspended in 750 μL Zymo BashingBead Buffer. The pellets were sonicated in a water bath sonicator for 15 min and then gently pipetted up and down. The skeleton fragments leftover from the MTBE:methanol extractions were added to the sample tubes and DNA extractions were performed using the Zymo Quick-DNA Fecal/Soil Microbe Miniprep Kit according to the manufacturer’s instructions. DNA extractions of whole coral tissue biopsies were performed using the same extraction kit and protocol. Symbiodiniaceae ITS2 sequences were amplified by qPCR using ITS-DINO and ITS2Rev2 primers [66, 67]. Relative amplicon yields were determined using the ΔΔCt method.

#### Proteomics

Similarly, we assessed the effect of peptide extraction method on coral proteomics profiles. Paired biopsies of *M. capitata* were either flash frozen in liquid nitrogen (n = 5) or fixed in acidified methanol (n = 4) and processed using a standard proteomics workflow or prior metabolite extraction using the described MTBE:methanol method, respectively. Briefly, flash frozen samples were suspended in 200 μL 4% SDC and immediately heated to 95 °C for 5 min to denature proteases and phosphatases. Samples were then probe-sonicated (2 x 10 s, 10% amplitude with resting on ice in between; Fisher Scientific Sonic Dismembrator Model 500) and centrifuged at 21,000 x g for 10 min to pellet any skeletal debris. The supernatant volumes were then readjusted to 200 μL, followed by sequential addition of 800 μL of methanol, 200 μL of chloroform, and 600 μL of water with thorough vortexing between each addition. Samples were centrifuged at 14,000 x g for 10 min, the supernatant removed, and the protein pellet washed twice with 400 μL of methanol and centrifugation at 20,000 x g. Cleaned pellets were dried for 5 min in a Speedvac (Labconco CentriVap SpeedVac System). Protein pellets obtained from the MTBE:methanol metabolite extraction were washed twice under identical conditions (400 μL of methanol, 20,000 x g centrifugation each time) and dried for 5 min in a Speedvac prior to downstream processing.

Subsequent steps were identical regardless of initial extraction method. Briefly, samples were resuspended in 200 uL of lysis buffer (8M urea in 0.1M Tris, 250mM NaCl, 50mM ammonium bicarbonate), and probe sonicated as before to resuspend the pellet. Samples were chemically reduced (10 mM TCEP, 30 min incubation at 23 °C) and alkylated (40 mM 2-CAM, 30 min incubation at 23 °C), then digested (1:50 Trypsin, 1:100 LysC, 16 h incubation at 23°C).

Subsequently, they were acidified, desalted using C18 tips (three washes with 0.1% TFA, elution by 50% ACN in 0.1% FA) and then dried. Samples were resuspended in 0.1% FA for proteomic analysis by LC-MS (see Supplementary Materials for detailed LC-MS method). Raw data were processed using Spectronaut (Biognosys, v20.3) [68]. Identified proteins were matched to human orthologs using OrthoFinder [69] and assigned gene ontology (GO) terms using the ‘*msigbr’* package in R. A rank-based GO enrichment was carried out using the ‘*GO_MWU’* package in R [70]. Effects of extraction methods were assessed by comparing presence/absence of unique proteins, differences in their abundances, and enrichment of cellular component GO terms (FDR-adjusted *p* < 0.05).

## Results and Discussion

Metabolomics provides a snapshot into the real-time physiological state of a coral, providing information on the structural composition of its cells, its energetic state, and its biotic and abiotic interactions by characterizing diverse compounds. This study aimed to provide guidelines and recommendations to facilitate the use of untargeted metabolomics by a wider research community interested in coral reefs. We have tried to simplify the approach to sample collection, highlight methodological limitations and provide new analytical tools to maximize the biological insight gained from coral holobiont samples.

### Challenges and considerations for coral biopsy collection for metabolomics analysis

#### Effects of metabolome quenching and sample handling methods

Field collection of coral samples often takes place in remote locations, where ideal sample preservation methods are not always feasible due to limited access to materials such as liquid nitrogen, delays in cold storage of samples, and the need to transport samples long distances for analysis. To capture the metabolome at a specific physiological state, cellular metabolism must be rapidly quenched. For tissue samples this is typically achieved by flash freezing in liquid nitrogen or immersion in cold or boiling solvents followed by immediate extraction or storage at -80 °C [71]. But the rapid turnover and high susceptibility of certain metabolites to enzymatic transformation make it difficult to preserve their native state even under ideal conditions [72]. Reliable metabolome fixation must therefore balance reproducibility, practicality, and field feasibility.

We assessed the effects of various metabolome quenching and sample handling methods on the metabolic profiles of four Scleractinian coral species, simulating optimal to sub-optimal scenarios of field sample collection, with the aim of understanding limitations in terms of resulting data quality (Figure 2). The methods tested included liquid nitrogen, ethanol, methanol, acidified methanol, boiling, and desiccation of methanol-fixed samples followed by 48-hour storage at room temperature (48h RT). Sample treatment had a significant but comparatively minor effect on metabolome composition (PERMANOVA R^2^ = 0.04, F = 2.82, *p* = 0.001), while biochemical differences among coral species remained dominant (R^2^ = 0.65, F = 80.82, *p* = 0.001; Figure S2). A significant treatment-by-species interaction (R^2^ = 0.06, F = 1.42, *p* = 0.001) indicated that species differed in sensitivity to fixation method: *M. capitata* was most sensitive (R^2^ = 0.35, F = 2.61, *p* = 0.004; Figure 2a), followed by *P. meandrina* (R^2^ = 0.31, F = 2.17, *p* = 0.023; Figure 2d), whereas *M. patula* (R^2^ = 0.23, F = 1.39, *p* = 0.14; Figure 2b) and *P. compressa* (R^2^ = 0.20, F= 1.20, *p* = 0.308; Figure 2c) were not significantly impacted (Figure 2).

**Figure 2.**
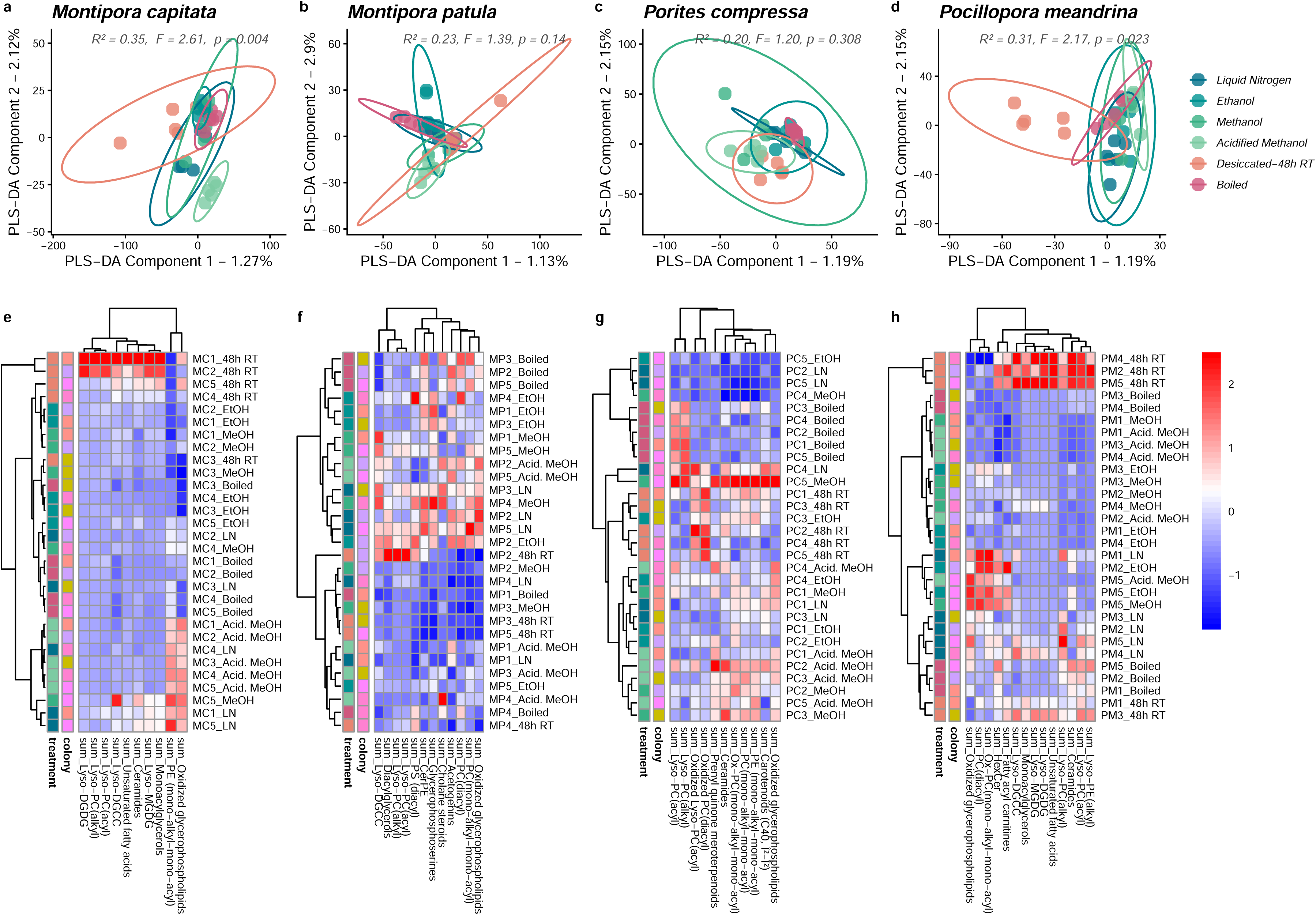
The effect of different metabolome quenching methods and sample handling on coral holobiont metabolic profiles. **(a-d)** Partial Least Squares Discriminate Analyses (PLS-DA) and PERMANOVA indicate differences in the profiles based on sample fixation in liquid nitrogen, ethanol, methanol, acidified methanol, by sample boiling, as well as the effect of sample desiccation for shipping at room temperature (48h RT). Results of PERMANOVA analysis are indicated within each plot. **(e-h)** Heatmaps of z-scaled summed abundance values of metabolite classes that were significantly clustered towards high PLS-DA component 1 and component 2 loading scores (Wilcox Rank Sum test, FDR-adjusted *p* < 0.05).

Metabolome profiles of *M. capitata* were most disturbed in ethanol, boiled, and 48h RT treatments, which differed compared to liquid nitrogen or acidified methanol fixed samples. Incubation of desiccated samples at room temperature resulted in increases in lyso-lipids and decreases in oxidized glycerophospholipids and ether-linked PE (Figure 2e). Colony-specific variation within *M. capitata* (notably samples associated with colonies ‘MC1’ and ‘MC2’) suggests a potential influence of symbiont type on the extent of sample degradation, as this host species commonly occurs with either *Durusdinium-* or *Cladocopium-*dominant symbiont communities in Kāneʻohe Bay [73]. Similar treatment effects were observed in *P. meandrina*, particularly elevated lyso-lipids and reduced oxidized glycerophospholipids and PC following desiccation (Figure 2h).

#### Effects of temporary storage duration

We next examined how delays between fixation and safe storage (in a -80 °C freezer) influence metabolomic profile stability (Figure 3). Following quenching in cold methanol, storage on ice (∼4 °C) for up to 24 h caused measurable degradation (PERMANOVA R^2^ = 0.04, F = 2.07, *p* = 0.017), characterized by increases in lyso-lipids and decreases in diacyl lipids such as PC, PE, and PS (Figure 3a). These changes likely reflect sustained activity of lipases. Species-specific responses were evident, with *Montipora* sp. being most, and *P. compressa* least, sensitive to degradation. Acidification of methanol with 1% formic acid effectively stabilized samples during prolonged storage on ice (PERMANOVA R^2^ = 0.02, F = 1.62, *p* = 0.076), preventing lyso-lipid accumulation (Figure 3a-c). In non-acidified methanol, an increase in lyso-PC only became significant after 24 h on ice for *M. capitata*, *P. compressa* and *P. meandrina* (Figure 3b), while lyso-DGDG rose significantly after just 1 h in *Montipora* samples, suggesting particularly high sensitivity of this lipid class to enzymatic derivatization in this genus (Figure 3c). The pH of the quenching solvent also had a significant impact on the signal intensity of PS irrespective of storage time (Figure 3e).

**Figure 3.**
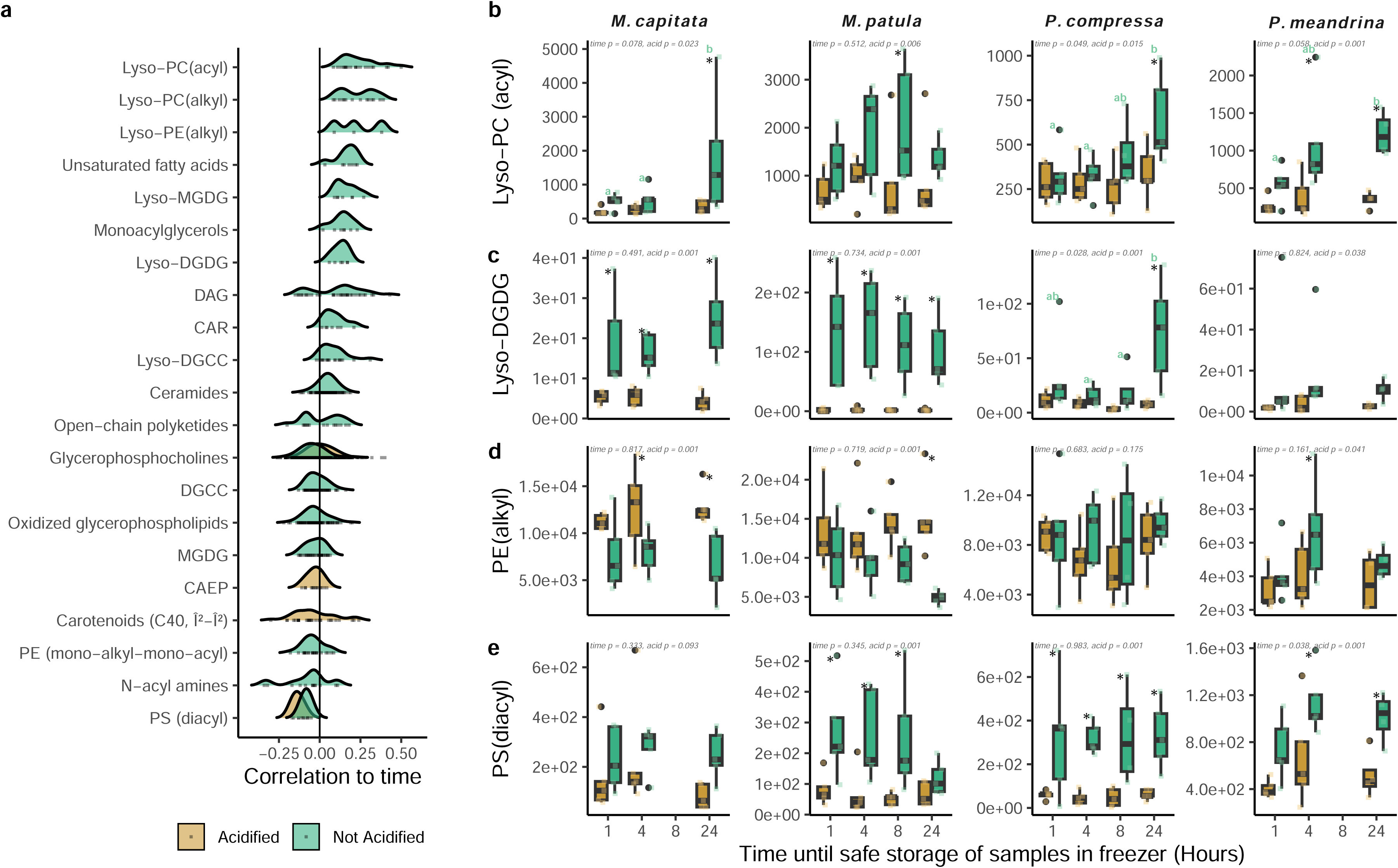
The effect of temporary storage duration (the time delay from sample collection until safe storage in a -80 °C freezer) on coral holobiont metabolomic profiles. **(a)** Metabolite classes with significant correlations to time among samples fixed in acidified or non-acidified methanol (Wilcox Rank Sum test, FDR-adjusted *p* < 0.05). **(b-e)** Effect of temporary storage duration on ice (∼4 °C) in acidified or non-acidified methanol on summed intensities of selected metabolite classes. Statistical significance for the main factors time and acidification were determined by 2-way ANOVA and indicated by the presence of an asterisk (significance of acidification) and the use of different letters (significance of time within acidified or non-acidified treatments).

#### Sampling recommendations based on test results

Although fixation method significantly influenced coral holobiont metabolomic profiles, the overall impact of using sub-optimal treatments was relatively modest and should not discourage researchers from conducting metabolomics analyses on field-collected corals. Instead, understanding the limitations of each method helps to avoid misinterpretation of results by accounting for metabolite susceptibility to enzymatic derivatization. Our analysis indicates that acidified methanol fixation is the most suitable approach for coral lipidomics, as it minimized lipase-derived artefacts and data variability (Figure 2). Methanol can be substituted with ethanol when the former is unavailable or restricted due to its higher toxicity and regulation. The addition of other additives such as antioxidants (*e.g.*, butylated-hydroxytoulene) could further prevent oxidation artifacts, which is particularly important for studies targeting oxidized lipids [74], but this was not tested here. Flash freezing in liquid nitrogen remains an excellent option if samples can remain frozen until extraction and analysis. The highest data variability resulted from the desiccation and 48 h RT treatment used to simulate transport of dry samples when cold shipping or solvent transport is restricted. While our method only involved drying by nitrogen gas, complete lyophilization is expected to substantially improve sample stability and is widely used for long-term storage and shipment of metabolomics samples. Based on these findings, we recommend that coral biopsies be quenched as rapidly as possible in acidified methanol, kept cold throughout collection (samples could optimally be stored on dry ice) and transferred to secure cold storage promptly. Short-term storage (up to several weeks) at −20 °C is acceptable, whereas −80 °C is recommended for long-term preservation of metabolite integrity [74].

#### Considerations of sample biomass

Immediate extraction of whole coral holobiont biopsies minimizes sample handling and processing time, preserving the native metabolome, and enabling high throughput sampling. A tradeoff, however, is that extracts cannot be normalized to conventional biomass metrics (*e.g.* total protein or symbiont density) prior to LC-MS analysis [37]. In the workflow presented here, sample biomass is relatively standardized by collecting fragments of a consistent size (5 mm diameter cores), and data are subsequently normalized using post-acquisition normalization procedures appropriate for untargeted metabolomics [75]. Nevertheless, inter- and intraspecific differences in tissue thickness can lead to variation in total biomass, and consequently, the maximal ion intensity measured. For instance, *Pocillopora acuta* exhibits notably thin tissue (Figure 4a; [76]). The relationship between maximal ion intensity and the number of detected features (*i.e.,* metabolite richness) follows a saturating curve, reflecting the finite linear dynamic range of the mass detector [77]. Once feature detection approaches saturation, variation in sample concentration no longer affects feature richness (Figure 4b). In contrast, severely reduced biomass – such as in heavily bleached corals – lowers metabolite concentrations and can result in substantially fewer detected features (Figure 4c), leading to variable metabolome coverage across samples. A comparable challenge is often addressed in genomics by data rarefaction [78], but no equivalent strategy has yet been broadly implemented for metabolomics data. Conversely, overly concentrated samples can exceed the linear dynamic range of the detector, leading to signal saturation, ion suppression and loss of quantitative linearity between signal intensity and metabolite abundance. Evaluating signal-to-noise ratios in pooled QC samples and adjusting sample concentrations or applying low-quality feature filtering can mitigate these issues [49]. Finally, when sample biomass is inherently limited – such as under sampling restrictions – alternative, minimally invasive techniques (*e.g.*, blunt-needle sampling of individual polyps) can still yield meaningful biochemical data [79].

**Figure 4.**
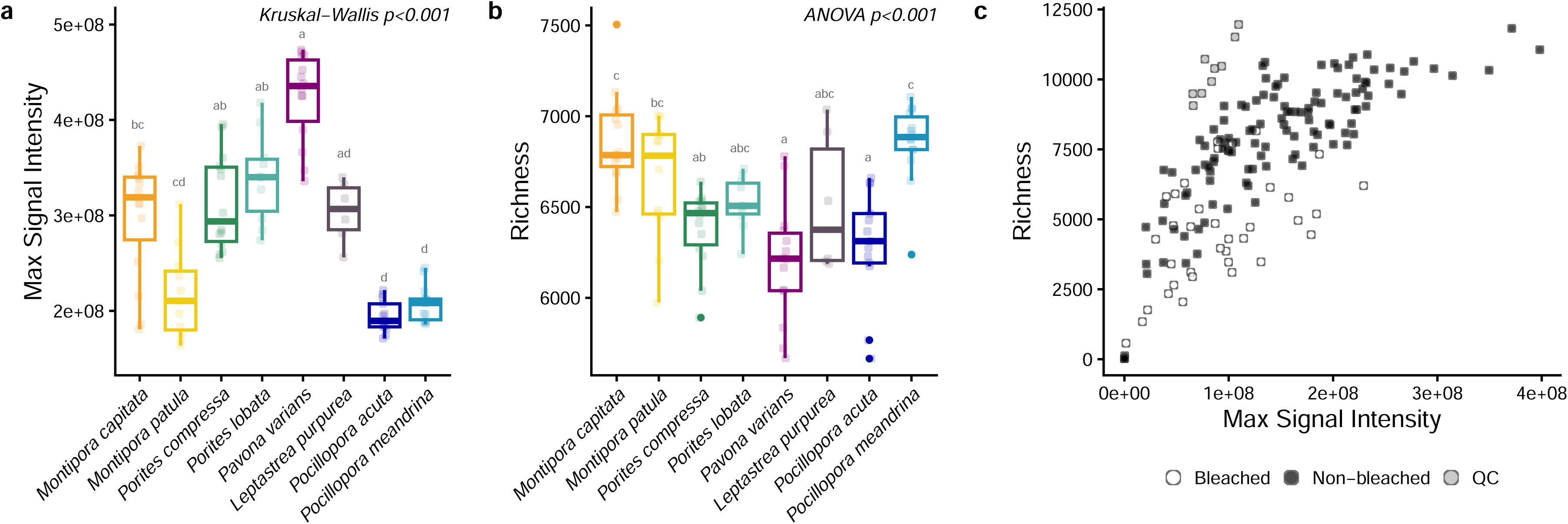
The correlations between sample concentration (maximal signal intensity) and the number of metabolite features detected (richness) in coral extracts. **(a)** The ion intensity of the feature with the highest signal intensity (a proxy for biomass) detected in distinct colonies of each of eight species of coral from Kāneʻohe Bay, Hawaiʻi, and **(b)** the total number of features detected (richness) in each of these samples after removal of non-biological features. **(c)** The relationship between maximal signal intensity and richness in a sample set of healthy and bleached corals from Curaçao.

### Accessible annotation of coral holobiont metabolomes

Metabolite annotation remains a major challenge in untargeted LC-MS/MS metabolomics [7, 10], with the large majority of detected features belonging to the uncharacterized “dark matter” of the metabolome [80]. The gold standard in annotation requires matching accurate mass (MS^1^), fragmentation spectrum (MS^2^), and retention time to an authentic analytical standard [81], but this is rarely achievable in untargeted analysis where thousands of features are detected, few of which have a standard commercially available. Annotation instead depends on matching spectra to compounds in public reference libraries [52].

Despite our relatively advanced knowledge of the composition of many lipid classes that constitute coral and Symbiodiniaceae lipidomes [17, 21, 24, 44, 59–63], this information has not previously been comprehensively compiled into an accessible format to enable automated annotation of coral spectra. To address this, we established a Coral Lipid Spectral Library, curated from a manually annotated dataset spanning multiple coral genera from Kāneʻohe Bay, Hawaiʻi, and several Symbiodiniaceae genera, both *in hospite* and in culture. This library includes 1,513 lipids, encompassing 18 classes. When applied to our dataset of four Hawaiian coral species used for testing sample fixation methods in the current study, the GNPS libraries provided 116 library matches (cosine score of >0.85). Using the Coral Lipid Library under the same criteria yielded 937 annotations. Combined, this increased the total to 968 annotated features out of 10,086 detected, raising the annotation rate from 1.15% to 9.6%. Similarly, the number of annotated features for our *G. fascicularis* dataset increased from 262 to 1532. While still low, even well-studied systems such as human serum or tissue samples typically only achieve around 10% annotation [10]. Further improvement in the coverage and accuracy of coral holobiont metabolome annotation will rely on community-driven efforts to expand and refine spectral libraries, enabling more reproducible and transparent metabolomic interpretation across studies.

### Holobiont metabolomics – Mapping metabolites to host and symbiont origin

Interpreting coral metabolomes requires knowledge of the organismal origin of metabolites – particularly, whether compounds are synthesized by the coral host or the algal symbiont. Generally, this is achieved by physically separating coral and algal fractions before analysis [17, 21, 41], which allows for better quantitation, and improved compound detection by reducing matrix complexity [37]. However, this approach is laborious, necessitates larger coral fragments, risks compromising the native metabolome due to increased sample handling and becomes impractical for large-scale studies. Extracting whole holobiont biopsies therefore offers a more practical alternative [16, 22, 82, 83] and can instead rely on analysis of carefully chosen biological reference samples related to the subjects, for example visually bleached corals with diminished symbiont biochemicals [30].

Certain compound classes have known coral- or algal-specificity. Algal-derived membrane lipid classes such as DGCC, MGDG, DGDG and SQDG occur exclusively in the algal symbiont [60, 62], whereas the coral host uniquely synthesizes the sphingophosphonolipid, CAEP [84]. Ether-linked lipids (*e.g.*, mono-alkyl-mono-acyl-PC, -PE, -PS and MADAG) are abundant in cnidarians [63, 85] but have not been detected in their algal symbionts [86]. The symbiotic partners also differ in the fatty acids they synthesize; for example, 18:4ω3 and 18:5ω3 are characteristic of symbiont galactolipids [87, 88], whereas 20:3ω6, 20:4ω6 and 22:4ω6 occur predominantly in coral host lipids [89]. However, most holobiont metabolites cannot be assigned to either partner based on chemistry alone.

We therefore developed a holobiont metabolomics framework to map metabolites to their coral host or algal symbiont source based on their presence/absence and relative abundance across representative coral and Symbiodiniaceae sample types in symbiotic and non-symbiotic or isolated states. Effect sizes from comparisons between algal cultures and bleached corals and between isolated algal pellets and healthy corals, were strongly correlated (slope = 1.17, R^2^ = 0.19, *p* < 0.001; Figure 5a), validating the integration of these four sample types for this purpose. Across >1000 holobiont samples representing eight Hawaiian coral species and >10,000 detected features, 64% of features could be mapped to the coral host (26.8%), algal symbiont (23.9%), or both (13.8%; Figure 5b). Thus, more than half of unannotated features were assigned a specific holobiont compartment, providing at least one level of information to the “dark matter” of the coral metabolome (Figure 5c). Unclassified features generally had low signal intensities, likely falling below the detection limit in isolated symbiont and bleached coral samples (Figure S5). Metabolite classes with high proportional classification as host metabolites included ether-linked PC, PE, PI and PS, ester-linked PS, lyso-PC and lyso-PE, ceramides, CerPE and CAEP (Figure 5c). Conversely, prominent algae-associated classes included DGCC, DGDG, carotenoids and apocarotenoids. The metabolite classes that were most common to both the algal and coral fraction included diacyl-PC, TAG, DAG, HexCer, Neutral glycosphingolipids and N-acyl amides (Figure 5c). While most glycerolipids (*e.g*. MAG, DAG, TAG) are common to both holobiont fractions, those with high proportional association with the coral host tissue relative to the algal symbiont contained 20:4, 22:4, 22:5, 18:0, 20:0, 22:0, 22:1, 16:4, and odd chain fatty acids (*e.g.* 19:0, 17:0, 21:0; Figure S4). Conversely, those with high proportional association with the algal symbiont relative to the coral contained characteristic algal PUFAs – 22:6, 18:4, 18:5, 18:3, 20:5, 28:8 – or shorter-chain saturated fatty acids – 16:0 and 14:0 (Figure S4).

**Figure 5.**
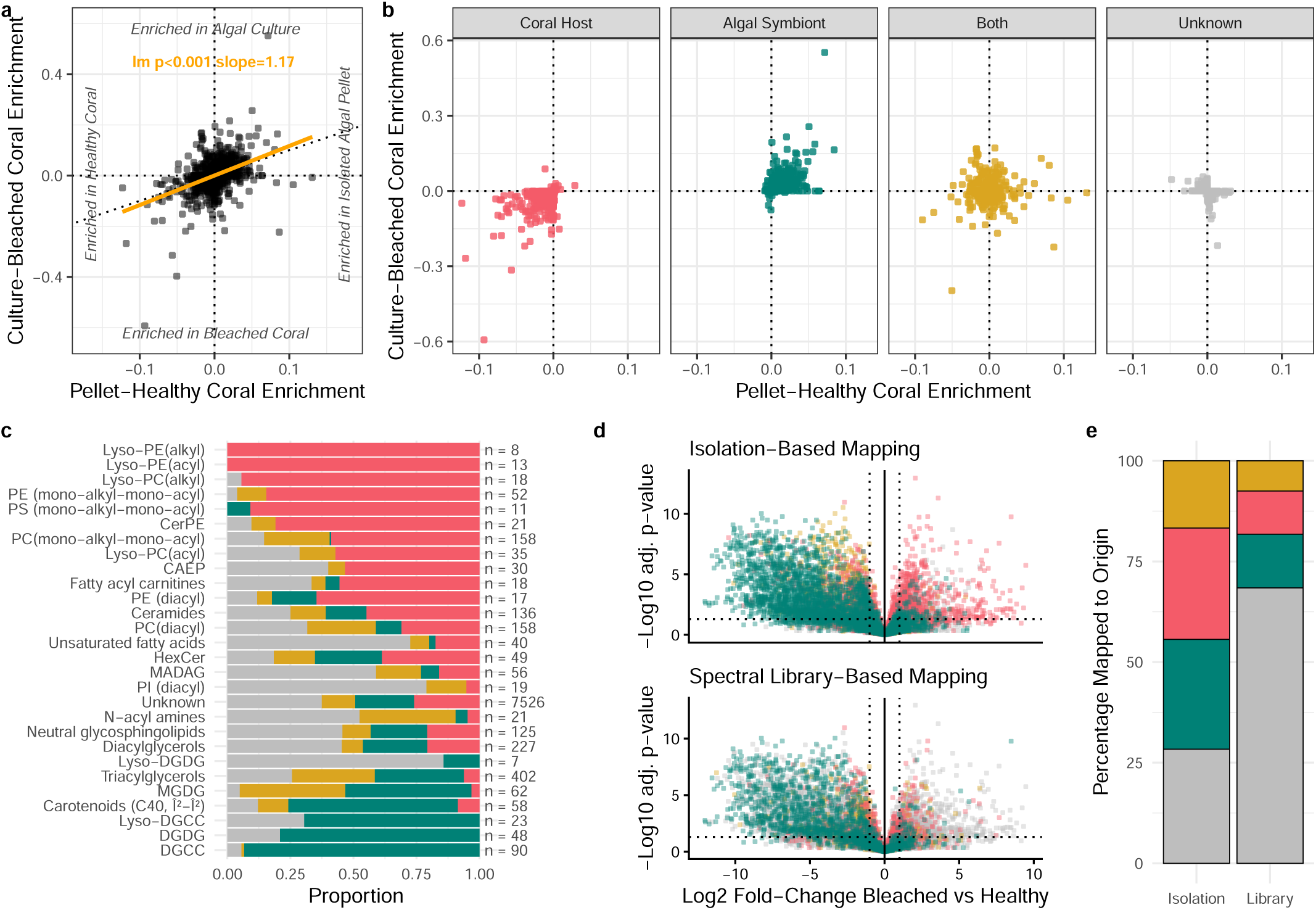
Mapping metabolites to host and symbiont origin. **(a)** Effect sizes of metabolite feature abundances derived from two separate generalized linear models, algal culture *versus* bleached coral and isolated algal pellet *versus* healthy coral. **(b)** Distribution of features mapped to coral host, algal symbiont, both host and symbiont, or unknown origin. **(c)** Proportions of organismal origin classification for selected lipid classes. **(d)** Origin mapping derived from internal sample or spectral library-based mapping imposed onto log2-fold change in metabolite abundance between chemically bleached and non-bleached *Galaxea fascicularis*. **(e)** Percentage of mapped features using isolation- or spectral library-based mapping.

Integrating organismal origin annotations into spectral libraries [90] could remove the need to run extracts of isolated coral and algal samples as part of every study, particularly when such sample types are inaccessible. To test this idea, we analyzed menthol-bleached and healthy *G. fascicularis* samples [91], and compared origin assignments derived from (1) spectral library matching to our curated reference dataset, and (2) mapping to isolated coral and algal extracts. Log_2_-fold changes revealed expected declines in algal-derived metabolites in menthol-bleached corals and more variable responses among coral compounds in response to bleaching (Figure 5d). Metabolite mapping to isolated reference samples classified 72% of features to coral host or algal symbiont origin, whereas the spectral library assigned 32%, yet with high accuracy (Figure 5e). Continued curation of spectral libraries will improve both coverage and reliability of origin assignments. Ultimately, this framework should be extended to integrate bacterial, fungal and viral metabolites to advance holistic biochemical profiling of coral holobionts, as these microbes also shape the coral metabolome [92, 93].

### Multi-omics from single samples to maximize data while minimizing destructive field sampling

Integrating metabolomics with transcriptome, proteome and microbiome profiles, can provide a more comprehensive understanding of biological systems [94], including coral reefs [95, 96]. For environmental samples, this typically requires separate specimens for metabolomics and the other data type [82], which is problematic given the magnitude of molecular differences that can occur over very small spatial scales [79]. Previous work has, however, shown that DNA and proteins can be preserved within metabolomic extraction protocols [45]. We evaluated whether DNA or protein could be co-extracted from single 5 mm coral biopsies for Symbiodiniaceae ITS2 amplification or proteomics, respectively. Sonication of biopsies in MTBE:methanol (3:1, v:v) containing 1% formic acid effectively extracted the coral tissue from the skeletal fragment, while the high solvent strength precipitated DNA and protein, allowing for separation from metabolites by centrifugation. We applied standard DNA and proteome extraction protocols to the resulting pellets and compared data quality to that of flash-frozen samples processed using dedicated extraction protocols. We note that, while we tested DNA and protein extractions on separate samples, these polymers could be isolated from single samples using classic protocols [97].

ITS2 amplification was successful as part of the metabolomics workflow and produced equivalent – or even higher in the case of *M. capitata* – relativized qPCR yield than dedicated DNA extraction (Figure 6a). Similarly, total protein counts were comparable between dual-extracted and dedicated proteomics workflows (Figure 6b), with substantial overlap in detected proteins (Figure 6c). Yet, metabolomics-derived proteomes exhibited differences in protein abundances (Figure 6d), with proteins uniquely identified in the metabolomics workflow being enriched in mitochondrial and organelle-linked complexes, including ribosomal subunits, suggesting preferential recovery of organelle-associated protein complexes (Figure 6e). Although our understanding of coral multi-omics data correlations remains sparse [98, 99], bioinformatics pipelines for integrating metabolomics data with other data types continue to advance. We advocate for broader adoption of multi-omics extraction strategies where feasible to maximize information from limited and valuable coral samples while reducing ecological impact and simplifying field collection.

**Figure 6.**
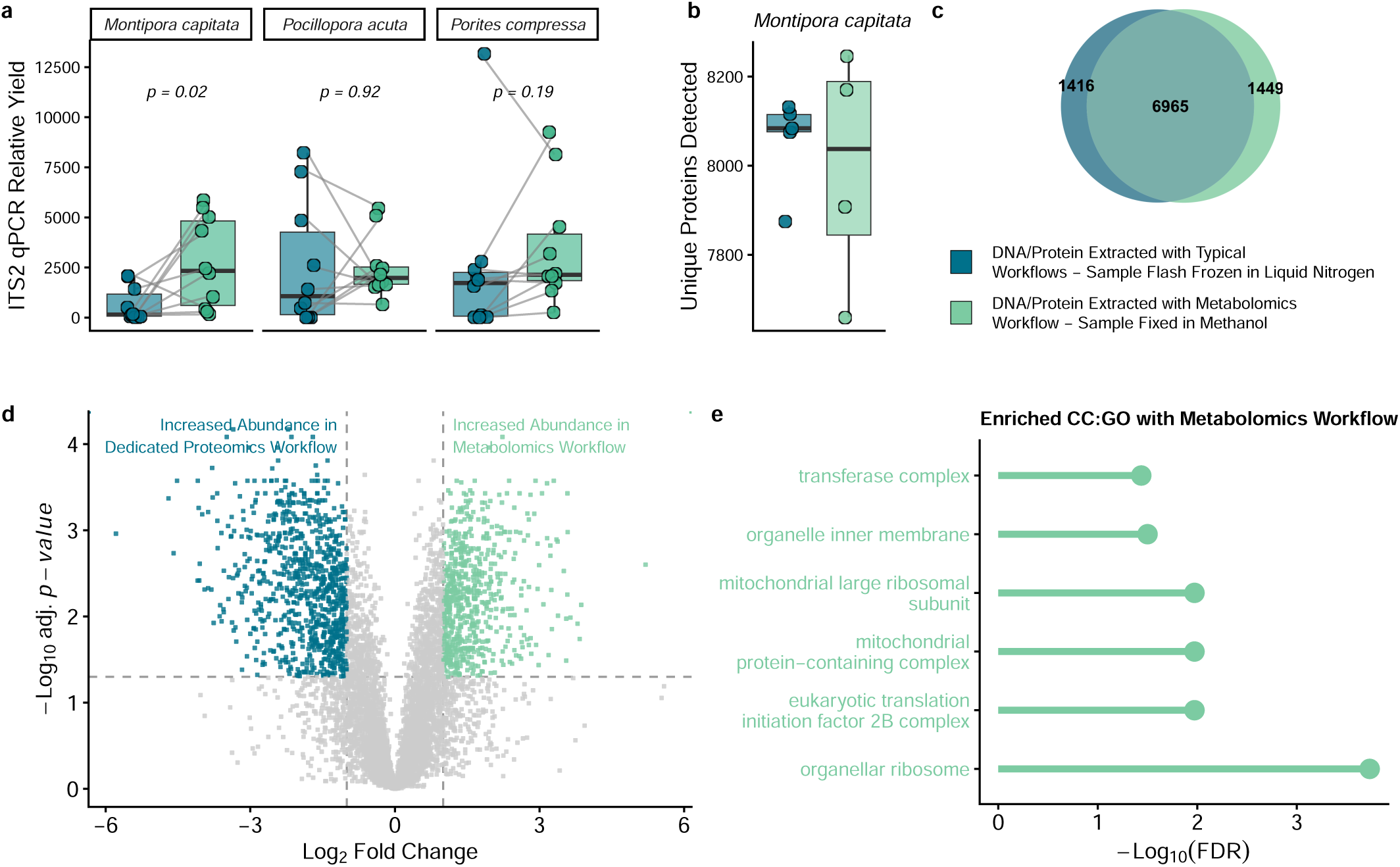
Comparisons of ITS2 amplification efficiency and coral proteomics profiles derived from typical DNA or protein extractions *versus* dual extractions from the metabolomics workflow. **(a)** Amplification efficiencies of algal symbiont ITS2 from *M. capitata*, *P. compressa*, and *P. acuta*, measured using qPCR relative yield of DNA either extracted from whole coral biopsies flash frozen in liquid nitrogen (blue) or from DNA extracted from precipitate pellets recovered following fixation of coral biopsies in methanol and metabolite extractions using MTBE:methanol (3:1, v:v; green). **(b)** The total numbers of unique *M. capitata* coral proteins identified using either a typical proteomics pipeline (proteins extracted from whole biopsies that were flash frozen in liquid nitrogen) or from dual metabolite/protein extraction using our optimal metabolomics workflow (biopsies fixed in acidified methanol). **(c)** Numbers of identified proteins that were common or unique between the two extraction methods. **(d)** Differential abundances of proteins identified in both extraction methods. **(e)** Enrichment of cellular components gene ontology terms in the fraction of uniquely identified proteins in the optimal metabolomics pipeline (note: there was no significant GO:CC enrichment in the uniquely identified proteins from the typical proteomics pipeline).

## Conclusion

The field of metabolomics is evolving rapidly and holds promise for advancing our understanding of complex biological systems. The simplified approach for sample collection, metabolite extraction and annotation of whole coral holobiont biopsies described here can facilitate larger-scale metabolomics analysis of corals across wider populations and geographic locations. Such high throughput studies could accelerate progress on pressing challenges such as coral thermal tolerance and disease susceptibility [100]. Our framework for mapping metabolites to holobiont sources is broadly applicable to other organisms and ecosystems. Continued commitment to open science and collaboration will be crucial for building a comprehensive, taxonomically-informed metabolome reference base. While access to specialized instrumentation for metabolomics analysis can be limited, continued efforts to forge global collaborations with mass spectrometry facilities or laboratories working in metabolomics should promote more inclusive scientific inquiry. Such collaborative efforts will deepen our understanding of microbial dynamics, community structure and metabolic processes, with critical implications for ecosystem health, sustainability and biodiversity in a rapidly changing world.

## List of abbreviations

BHT: butylated hydroxytoluene
CAEP: ceramideaminoethylphosphonates
CerPE: ceramide phosphoethanolamines
CerPI: ceramide phosphoinositols
DAG: diacylglycerols
DGCC: diacylglyceryl carboxyhydroxymethylcholines
DGDG: digalactosyldiacylglycerols
DGTS/A: diacylglyceryl trimethyl-homoserine/alanine
GC-MS: gas-chromatography–mass spectrometry
HexCer: hexosylceramides
HIMB: Hawaiʻi Institute of Marine Biology
HPLC: high-pressure liquid chromatography
ITS2: internal transcribed spacer 2
LC-MS/MS: liquid-chromatography–tandem mass spectrometry
MADAG: monoalkyldiacylglycerols
MAG: monoacylglycerols
MGDG: monogalactosyldiacylglycerols
MS1: precursor mass
MS2: fragmentation spectrum
MTBE: methyl-tert-butyl-ether
NL: neutral loss mass
P: product mass
PC: phosphocholines
PE: phosphoethanolamines
PI: phosphoinositols
PQN: probabilistic quotient normalization
PS: phosphoserines
QC: quality control
RT: retention time
SM: sphingomyelins
SQDG: sulfoquinovosyldiacylglycerols
TAG: triacylglycerols

## Acknowledgements

This work was funded by the Michigan State University Climate Change Research Support Program, the National Science Foundation Organismal Response to Climate Change Grant (NSF-ORCC-2307516) and the Defense Advanced Research Projects Agency Award (DARPA-HR001122C0134). This is HIMB contribution xx and SOEST contribution xx. We would like to acknowledge Christoph Benning at Michigan State University for advice during method development, and Stanley Lio at HIMB for photographs of coral biopsy extraction. We further acknowledge assistance from Dr. Maren Ziegler and the staff at Preuss Pets, Lansing, MI for assistance with coral rearing in Michigan.

## Author Contributions

SLR: designed and performed experiments, collected samples, data analysis, wrote the paper; TBC: designed and performed experiments, collected samples, data analysis, wrote the paper; KSh: performed experiments, data analysis; IA: collected samples, performed experiments, data analysis; KSm: performed experiments, collected samples; GT: performed experiments, collected samples; LKW: performed experiments, collected samples; JEP: designed experiments, collected samples; MB: designed experiments; CD: designed and performed experiments, collected samples; data analysis, funded the work; TNFR: designed and performed experiments, collected samples; wrote the paper, funded the work; RAQ: designed and performed experiments, collected samples, wrote the paper, funded the work; all authors edited and approved the final manuscript.

## Data availability

The spectra and feature quantification files of all datasets analyzed during the current study are available in the Mass Spectrometry Interactive Virtual Environment (MassIVE) repository under the accession number MSV000100504.

