## Supplementary Materials for "From SCUBA to spectra: Broadly applicable methods for coral metabolomics research"

*Corresponding authors

[†]Detailed author list in supplementary file

### Metabolomics analysis of coral holobiont biopsies – Detailed protocol

### Step 1 – Coral biopsy collection and storage

#### Materials and Equipment needed:

- 5 mm dermal curette (*e.g.* for massive coral species)
- Bone clippers/wire cutters (*e.g.* for branching species)
- Forceps for handling small samples
- 1.5 mL amber glass vials with caps (1 vial per sample)
- HPLC-grade methanol
- LC-grade formic acid (to prevent lipase activity during sample fixation)
- butylated hydroxytoluene (BHT; acts as antioxidant to prevent artificial metabolite oxidation)
- 50 mL tube filled with 70% methanol
- 50 mL tube filled with MQ water
- Insulated box with ice

#### Protocol:

1.1. Label one 1.5 mL glass vial for each sample that will be collected using a unique sample ID.

1.2. Make a 10% BHT working solution in HPLC-grade methanol (w:v; using glass container)

1.3. Make up a stock of HPLC-grade methanol containing 1% formic acid and 0.01% BHT using glassware cleaned for LC-grade standard.

1.4. In the lab, pre-fill each 1.5 mL vial with 500 µL HPLC-grade methanol containing 1% formic acid and 0.01% BHT. Ensure the cap is sealed tightly to prevent evaporation.

1.5. Pre-chill the vials containing methanol prior to sample collection by storing at -20°C.

1.6. Prepare two 50 mL tubes that will be used for washing sampling instruments during collection. One tube should be filled with 70% methanol and one with purified water.

1.7. During sample collection, keep the glass vials containing methanol on ice in an insulated box (optionally, dry ice could be used to keep samples cold if sample collection is expected to take longer than ~8 hours).

1.8. Collect a small biopsy of coral either by pressing and twisting a 5 mm dermal curette into a massive coral to extract a tiny core of tissue and skeleton (for example for Porites), or by clipping a small piece or coral using bone clippers or a similar tool (for example for Pocillopora) (see Figure S1). Using forceps, quickly blot the biopsy onto tissue paper to remove excess seawater, then transfer the sample into a 1.5 mL glass vial containing acidified methanol. The biopsies can either be collected directly from corals in the field or aquaria, or else larger fragments can be collected and bought back to the boat or land in zip lock bags, where the smaller biopsies can be sampled and transferred to the vials. Optimally, 3-5 technical replicate samples should be collected for each coral colony. Ensure caps are sealed tightly to prevent evaporation.

1.9. Samples should remain cold and be transferred to a -80°C freezer as soon as possible.

**Note**: For sampling of coral for metabolomics analysis, the aim should be to preserve the coral as quickly as possible to prevent degradation or alteration of the metabolome. The protocol described above is one option, whereby the samples are fixed in acidified methanol. Other options for samples fixation such as flash freezing in liquid nitrogen are described in the main manuscript.

### Step 2 – Metabolite extraction

**Materials and Equipment needed:**

- HPLC-grade methyl tertiary butyl ether (MTBE) 100%
- HPLC-grade Methanol 100%
- HPLC-grade or MQ water
- LC-MS-grade formic acid
- butylated hydroxytoluene (BHT; 10% w:v in methanol)
- 1 x 1.5 mL microcentrifuge tube labelled with sample ID and "DNA/protein" per sample
- 1 x 1.5 mL microcentrifuge tube labelled with sample ID to use for phase separation step per sample
- 1 x 1.5 mL microcentrifuge tube labelled with sample ID and "polar" per sample
- 1 x 1.5 mL amber glass vial labelled with sample ID and "lipid" per sample
- Vortex
- Water bath ultrasound sonicator
- Desktop centrifuge
- Nitrogen gas sample evaporator

**Note:** All glassware used should be dedicated for metabolomics. Glass measuring cylinders and bottles should be rinsed 2-3 times with water (HPLC-grade or MQ) and each of the solvents to be used (HPLC-grade). All plastics used with solvents (microcentrifuge tubes, pipette tips) should be high-grade for to solvent resistance to reduce plastic polymer contamination.

The metabolite extraction method is adapted from Salem et al. [1].

**Protocol:**

2.1. Prepare solvents needed for extractions. Make up as much solvent as is needed to extract an experimental sample set so that a single fresh batch of solvents can be used for all samples:

- MTBE:Methanol 3:1 v:v with 1% formic acid and 0.01% BHT (e.g. to make up 20 mL: 15 mL MTBE + 5 mL methanol + 200 μL formic acid + 20 μL 10% BHT)
- Water:methanol 3:1 v:v (e.g. to make up 20 mL: 15 mL water + 5 mL methanol)

2.2. Pre-chill solvents at -20°C

2.3. Get the coral biopsies out of the -80°C freezer and place on ice

2.4. If the coral biopsies are fixed in methanol, evaporate the methanol using a Nitrogen gas evaporator (option: if you are confident that methanol volumes are consistent – *i.e.* no methanol evaporated or spilled during sample collection and storage – you can skip this step and instead add an appropriate volume of MTBE containing formic acid and BHT to make up to the necessary solvent ratios in step 2.5.).

2.5. Add 500 μL ice-cold MTBE:methanol (3:1, v:v, 1% formic acid + 0.01% BHT) to each vial.

2.6. Vortex each vial thoroughly (10-20 seconds)

2.7. Sonicate each sample in a water bath sonicator for 15 minutes (glass vials can be held in a foam sample holder to float in the water bath during sonication. Make sure water from the water bath cannot enter sample). Add ice to the water in the water bath sonicator to keep samples cold during sonication.

2.8. Vortex each vial thoroughly again (10-20 seconds)

2.9. Transfer the full 500 μL of solvent into a 1.5 mL microcentrifuge tube labelled “DNA/protein”, leaving the coral skeleton fragment behind.

2.10. Centrifuge at 15’000 x g for 10 minutes to pellet protein/DNA/debris (centrifuge at 4°C if possible)

2.11. Transfer the supernatant into a new 1.5 mL microcentrifuge tube for phase separation

2.12. The precipitate pellet from step 2.10 can be preserved and used for protein and/or DNA extraction for proteomics and ITS2 sequence analysis, respectively. To preserve the pellet, wash it once with 500 µL 100% methanol by centrifugation (15’000 x g, 10 mins) without disturbing the pellet to remove any remaining MTBE. Pellets can then be stored in methanol at -80°C. To proceed with DNA or protein extraction, remove all methanol and allow the pellet to air dry before re-suspending in appropriate extraction buffer. Sonicate in a water bath sonicator for 10 min and pipette up and down gently to re-suspend precipitate. For DNA extraction, the leftover skeleton fragment from step 2.11 can be added to the re-suspended precipitate. Proceed with typical DNA and protein extraction protocols.

2.13. To each phase separation tube from step 2.11, add 500 µL ice-cold water:methanol (3:1, v:v)

2.14. Vortex each vial thoroughly (10-20 seconds)

2.15. Incubate at -20°C for 30 min for phase separation

2.16. Centrifuge at 10’000 x g for 10 minutes (centrifuge at 4°C if possible)

2.17. Carefully transfer the upper organic phase into a 1.5 mL glass vial labelled “Lipid”. Avoid transferring any of the lower aqueous phase. Transferring 2 x 120 µL should collect most of the organic phase.

2.18. Collect the bottom aqueous phase into a 1.5 mL microcentrifuge tube labelled “polar”. Transferring 650 µL in one go works well to collect most of the aqueous phase.

2.19. Dry the extracts under a stream of nitrogen gas

2.20. Dried extracts can be stored at -80°C until analysis.

**Note:** While performing extractions of a set of samples you should always include 2-3 extraction blanks as these will be required to identify and remove features in the data that are derived from non-biological factors encountered during the extraction such as plastics.

### Step 3 – Data acquisition by LC-MS/MS – Reverse Phase method for Lipids

**Materials and Equipment needed:**

- High resolution mass spectrometer such as Thermo Q-Exactive
- High-pressure liquid chromatography system such as Thermo Vanquish
- C18 reverse phase HPLC column (Acquity UPLC BEH C18 column (130 Å, 1.7μm, 2.1 x 100 mm, Waters) (optional with replaceable guard column to increase lifetime of your column - VanGuard pre-column (Acquity UPLC BEH C18, 130 Å, 1.7μm, 2.1 x 5 mm, Waters))
- 96-well plate with cover slip or glass vials with inserts and caps
- HPLC-grade water
- HPLC-grade 2-propanol
- HPLC-grade acetonitrile
- LC-grade formic acid
- Ammonium formate

**Protocol:**

3.1. Make up the resuspension solvent for "Lipid" samples: 2-propanol : acetonitrile : water (2:1:1 v:v:v) and pre-chill at -20°C.

3.2. Add 500 μL lipid resuspension solvent to each sample and vortex.

3.3. Prepare a quality control "QC" sample by transferring around 20 μL of each sample in your experimental sample set to a 1.5 mL glass vial (adjust volume depending on how much total volume of QC needed). This pooled sample should be run at regular intervals to ensure consistent performance of the chromatography and mass spectrometer and for downstream post-acquisition data normalization.

3.4. Transfer around 80 μL of each sample either into a 96-well plate or into a vial with a glass insert. Samples should be arranged in a randomized order with respect to treatment groups. A solvent blank and a QC sample should be acquired once per every ten samples.

3.5. Optional: Make up a serial dilution of the QC, consisting of 8-10 concentrations with 2-3 technical replicates, and add to sampler. This is optional but recommended for post-acquisition data normalization and for downstream high quality feature filtering.

3.6. Make up the LC running solvents. Especially solvent B should be made up fresh.

- **Solvent A**: acetonitrile : water (60:40, v:v) containing 1% formic acid and 10mM ammonium formate
  For 1L of solvent A, weigh out 0.63 g of ammonium formate and dissolve in 1 L glass bottle with 400 mL HPLC-grade or MQ water. Add 1 mL LC-grade formic acid. Add 600 mL HPLC-grade acetonitrile. Mix well by swirling.
- **Solvent B**: 2-propanol : acetonitrile (90:10, v:v) containing 1% formic acid and 10mM ammonium formate
  For 1L of solvent B, weigh our 0.63g of ammonium formate and dissolve in 1 mL of HPLC-grade water in a small glass beaker before transferring to a 1 L glass bottle. Add 900 mL of HPLC-grade 2-propanol. Add 1 mL of LC-grade formic acid. Mix well by swirling. Add 100 mL of HPLC-grade acetonitrile. Note: ammonium formate may come out of solution. Make solvent B the day before running to let it dissolve fully overnight.

3.7. Attach solvents to LC system and purge solvent lines.

3.8. Attach column to LC system and set column compartment temperature to 55°C.

3.9. Add samples to sampler compartment and set temperature to 15°C.

3.10. Setup LC and mass spectrometer method parameters (see tables below)

3.11. Set the flow rate to 0.4 mL/min and check that the pressure stabilizes.

3.12. Run a couple of blanks and QC samples to check method performance and data quality.

3.13. Optional: run extraction blank and put major peaks into the exclusion list in the mass spectrometer method setup.

3.14. Set sample sequence to run using 10 uL injection volumes and acquire data using data-dependent acquisition (DDA). Data can be acquired in positive and negative ionization modes, but priority should be placed on acquiring the data in positive ionization mode.

**Table S1: Chromatography gradient for lipidomics method**

| Time (min) | Flow rate (mL/min) | % B |
|---|---|---|
| 0.0 | 0.4 | 20 |
| 1.0 | 0.4 | 20 |
| 2.0 | 0.4 | 43 |
| 10.0 | 0.4 | 55 |
| 10.1 | 0.4 | 65 |
| 17.0 | 0.4 | 90 |
| 17.1 | 0.4 | 99 |
| 18.0 | 0.4 | 99 |
| 18.1 | 0.4 | 20 |
| 20.0 | 0.4 | 20 |

**Table S2: Mass spectrometer parameters for lipidomics method**

| | |
|---|---|
| **Full MS** | |
| **Resolution** | 35,000 |
| **AGC target** | 3e6 |
| **Max IT** | 100 ms |
| **Scan range** | 200 – 1500 m/z |
| **dd-MS2** | |
| **Resolution** | 17,500 |
| **AGC target** | 1e5 |
| **Max IT** | 50 ms |
| **Loop count** | 5 |
| **Top N** | 5 |
| **Isolation window** | 1 m/z |
| **Stepped NCE** | 20, 40, 60 |
| **dd Settings** | |
| **Min AGC target** | 5e3 |
| **Intensity threshold** | 1e5 |
| **Dynamic exclusion** | 3s |

### Step 4 – Data acquisition by LC-MS/MS – HILIC method for Polar metabolites

**Materials and Equipment needed:**

- High resolution mass spectrometer such as Thermo Q-Exactive
- High-performance liquid chromatography system such as Thermo Vanquish
- HILIC HPLC column (Acquity Premier BEH amide VanGuard fit column (1.7μm, 2.1 x 100 mm, Waters)
- HPLC-grade water
- HPLC-grade acetonitrile
- LC-grade ammonium hydroxide
- Ammonium acetate

**Protocol:**

4.1. Make up the resuspension solvent for "polar" samples: acetonitrile : water (90:10 v:v; use aliquot taken from LC running solvent B) and pre-chill at -20°C.

4.2. Add 500 μL of polar resuspension solvent to each sample and vortex.

4.3. Prepare a quality control "QC" sample by transferring around 20 μL of each sample in your experimental sample set to a 1.5 mL glass vial (adjust volume depending on how much total volume of QC needed). This pooled sample should be run at regular intervals to ensure consistent performance of the chromatography and mass spectrometer and for downstream post-acquisition data normalization.

4.4. Transfer around 80 μL of each sample either into a 96-well plate or into a vial with a glass insert. Samples should be arranged in a randomized order with respect to treatment groups. A solvent blank and a QC sample should be acquired once per every ten samples.

4.5. Make up a serial dilution of the QC, consisting of 8-10 concentrations with 2-3 technical replicates, and add to sampler. This is optional but recommended for post-acquisition data normalization and for downstream high quality feature filtering.

4.6. Make up the LC running solvents. Especially solvent B should be made up fresh.
**Solvent A**: water with 10 mM ammonium acetate and 0.1% ammonium hydroxide
For 1 L of solvent, weigh out 0.77 g of ammonium acetate and dissolve in 1 L glass bottle with 1 L water (HPLC-grade or MQ). Add 1ml ammonium hydroxide. Mix well by swirling.

**Solvent B**: acetonitrile : water (90:10, v:v) with 10mM ammonium acetate and 0.1% ammonium hydroxide

For 1 L of solvent, weigh our 0.77 g of ammonium acetate and dissolve in 1 L glass bottle with 100 mL water (HPLC-grade or MQ). Add 1 mL ammonium hydroxide. Add 900 mL HPLC-grade acetonitrile. Mix well by swirling.

4.7. Attach solvents to LC system and purge solvent lines.

4.8. Attach column to LC system and set column compartment temperature to 40°C.

4.9. Add samples to sampler compartment and set temperature to 10°C.

4.10. Setup LC and mass spectrometer method parameters (see tables below)

4.11. Set the flow rate to 0.4 mL/min and check that the pressure stabilizes.

4.12. Run a couple of blanks and QC samples to check method performance and data quality.

4.13. Optional: run extraction blank and put major peaks into the exclusion list in the mass spectrometer method setup.

4.14. Set sample sequence to run using 10 uL injection volumes and acquire data using data-dependent acquisition (DDA). Data can be acquired in positive and negative ionization modes, but priority should be placed on acquiring the data in positive ionization mode.

**Table S3: Chromatography gradient for polar metabolites method**

| **Time (min)** | **Flow rate (mL/min)** | **% B** |
|---|---|---|
| 0.0 | 0.4 | 99 |
| 1.0 | 0.4 | 99 |
| 2.0 | 0.4 | 70 |
| 3.5 | 0.4 | 60 |
| 4.0 | 0.4 | 60 |
| 4.1 | 0.4 | 99 |
| 10.0 | 0.4 | 99 |

**Table S4: Mass spectrometer parameters for polar metabolites method**

| | |
|---|---|
| **Full MS** | |
| **Resolution** | 35,000 |
| **AGC target** | 1e6 |
| **Max IT** | 100 ms |
| **Scan range** | 70 – 1000 m/z |
| **dd-MS2** | |
| **Resolution** | 17,500 |
| **AGC target** | 1e5 |
| **Max IT** | 50 ms |
| **Loop count** | 5 |
| **Top N** | 5 |
| **Isolation window** | 1 m/z |
| **Stepped NCE** | 20, 40, 60 |
| **dd Settings** | |
| **Min AGC target** | 5e3 |
| **Intensity threshold** | 1e5 |
| **Dynamic exclusion** | 3s |

### Supplementary Methods

#### Data processing parameters

All LC-MS/MS datasets were processed in MZmine software (v 4.3) [2]. The specific parameters used for feature detection and alignment of the dataset used for metabolome quenching and sample handling method testing are indicated in table S5. This dataset encompassed biopsies of four coral species (*M. capitata*, *M. patula*, *P. compressa*, *P. meandrina*). After processing, feature quantification tables and MS/MS spectral files were exported using the export option for molecular networking (GNPS, FMBN). Feature-based molecular networking [3] was performed using $MS^1$ and $MS^2$ mass errors of 0.02, and a minimum of 0.65 cosine score and 4 matched peaks.

**Table S5**: MZmine processing parameters

| Processing step | Specific parameter | Lipid (ES+) |
|---|---|---|
| **Mass detection** | MS1 noise level | 4.0E4 |
| | MS2 noise level | 8.0E3 |
| **Chromatogram builder (ADAP)** | Min. consecutive scans | 5 |
| | Min. intensity | 4.0E4 |
| | Min. absolute height | 1.6E5 |
| | mz tolerance | 0.006 or 10ppm |
| **Chromatogram resolving (local minimum resolver)** | Chromatographic threshold | 95% |
| | Min. search range | 0.06 |
| | Min. relative height | 1.5 |
| | Min. absolute height | 1.6E5 |
| | Min. ratio of peak | 1.25 |
| | Peak duration | 0.06-1.25 |
| | Min. scans | 5 |
| **13C Isotope filter** | mz tolerance | 0.003 or 5ppm |
| | RT tolerance | 0.1 |
| | Max charge | 2 |
| **Alignment (Join Aligner)** | mz tolerance | 0.006 or 10ppm |
| | RT tolerance | 0.2 |
| | weight for mz | 75 |
| | weight for RT | 25 |
| **Feature list row filter** | Min samples | 4 or 1% |
| | Min features in isotope pattern | 3 |
| **Gap filling** | intensity tolerance | 20% |
| | mz tolerance | 0.004 or 7.5ppm |
| | RT tolerance | 0.15 |
| | Min. scans | 5 |

| | | |
|---|---|---|
| **Duplicate feature filter** | Filter mode | Old Average |
| | mz tolerance | 0.004 or 7.5ppm |
| | RT tolerance | 0.15 |

**Isolation of *in hospite* algal symbionts for metabolite mapping**

We separated algal symbionts from coral host tissue to obtain samples of symbiotic algae depleted of coral host tissue. This type of sample was used for mapping metabolites to coral host and algal symbiont compartments. Algal symbionts were isolated from five colonies of each of *Montipora capitata*, *Montipora patula*, *Porites compressa*, *Porites lobata*, *Pavona varians*, *Leptastrea purpurea, Pocillopora acuta and Pocillopora meandrina*. All coral were maintained in flow-through tanks at the Hawai‘i Institute of Marine Biology. 2-4 cm fragments were collected from each colony and put into individual zip lock bags. We added 1.5 mL of filtered seawater to each bag and removed coral tissue from the skeleton by blasting the fragment using an airbrush attached to a SCUBA tank. The tissue slurry was then transferred to a 1.5 mL microcentrifuge tube, placed on ice, and passed through a 12-gauge syringe needle ten times. We then performed 5 rounds of centrifugation at 500 x g for 5 mins to wash the algal symbiont pellet using 1 x PBS. Clean symbiont pellets were stored at -80°C until being extracted using the described metabolite extraction protocol.

***Galaxea fascicularis* menthol bleaching experiment**

One liter of filtered artificial seawater (FASW - 0.22 µm) was added to each of four 5 L plastic containers held in a water bath maintained at 26 °C. A submersible pump was added to each container to ensure a constant flow. Light was provided by LED lamps on a 12:12 h light:dark cycle. Replicate fragments obtained from two *Galaxea fascicularis* genotypes, hosting either *Durusdinium* or *Cladocopium* symbionts, were distributed into separate containers approximately two hours after the beginning of the light period. Coral fragments associated with each symbiont-type were distributed among two containers – control and treatment – with eight replicates per container. FASW of the experimental treatment containers was supplemented with 20% w/v menthol in ethanol to a concentration of 0.38 mM menthol [4]. The treatment containers were covered with black electrical tape to prevent light from entering, and a lid was fixed tightly to prevent menthol evaporation. After eight hours, the colonies were removed and temporarily submerged in fresh FASW in separate containers while the experimental and control containers were washed and refilled with 1 L of fresh FASW without menthol. Colonies were returned to their respective containers overnight. This treatment procedure was repeated for 5 consecutive days, followed by 2 days without menthol treatment or water changes. All colonies were fed freshly hatched artemia on the second and fifth days of treatment. A total of 20 days of menthol exposure was required before colonies became visibly bleached. Whole colonies were then sampled into 1.5 mL microcentrifuge tubes and flash frozen in liquid nitrogen before being stored at -80 °C. Samples were extracted and analyzed by LC-MS/MS as described in the main manuscript.

**Mass spectrometry proteomics acquisition and data processing**

Dried peptides were resuspended in 0.1% (v/v) formic acid in MS-grade water (Fisher Scientific) and analyzed on a timsTOF HT mass spectrometer, paired with a Vanquish Neo UHPLC system. Mobile phase A consisted of 0.1% (v/v) formic acid in MS-grade water (Fisher Scientific), and mobile phase B consisted of 0.1% (v/v) formic acid in 100% MS-grade acetonitrile (Fisher Scientific). The LC was operated in trap-and-elute mode, where the peptides were first trapped onto a PepMap Neo Trap column (5 mm, 100 Å pore size, 5 µm particle size) and then reversed-phase separated using gradients mentioned below on an Aurora Elite C18 reverse phase column (15 cm, 100 Å pore size, 1.5 µm particle size for captive spray, IonOptiks), kept at 50°C using a column oven for Bruker Captive Spray source (Sonation Lab Solutions), and ionized in a CaptiveSpray source (Bruker Daltonics) at 1700 V. For global proteome analysis, the %B gradient used was: 5% to 35% over 37 min at 0.3 µL/min, then to 45% in the next 4 mins, and 60% in the next 1 min, followed by an increase to 95% B over 3 min. All MS data was acquired in DIA-PASEF mode with variable isolation window widths in the m/z vs ion mobility plane. These windows were adjusted to maximize the coverage of precursor ions. MS1 scans were acquired from 100–1700 m/z and a dual-TIMS analyzer ramp rate of 9.42 Hz, with 100 ms accumulation and ramp times (100% duty cycle). The ion mobility range was 0.65–1.41 V·s/cm², isolation windows were 25 Da (1 Da overlap) with 46 mass steps per 1.27 s cycle, and collision energy decreased linearly from 59 eV at $1/K_0$ = 1.41 V·s/cm² to 20 eV at $1/K_0$ = 0.65 V·s/cm², collecting MS/MS spectra from 292.4–1397.4 m/z.

The raw files were processed with Spectronaut (Biognosys) with the directDIA+ (Deep) search algorithm, its in silico-derived DIA analysis. Carbamidomethylation (cysteine) was set as a fixed modification for database search. Acetylation (protein N-term) and oxidation (methionine) were set as variable modifications. The third version of the Hawaiian *Montipora capitata* proteome (http://cyanophora.rutgers.edu/montipora/) file was used for spectral matching. The false discovery rates for the PSM, peptide, and protein groups were set to 0.01. For MS2-level area-based quantification, the cross-run normalization option was unchecked (normalization was performed later using MSstats). Quantitative analysis was performed in the R statistical programming language (v.4.4.1). Initial quality control analyses, including inter-run clustering, correlations, principal component analysis (PCA), peptide and protein counts, and intensities were completed in R. Statistical analysis of protein abundance changes between exposed and control samples were computed using the R package MSstats (v 4.16.1). All peptides mapping to the same proteins were summarized together using the Tukey's Median Polish approach. MSstats performs normalization by median equalization, imputation was turned Off (set to FALSE), and statistical tests of differences in intensity between conditions was calculated using default settings in MSstats. Specifically, MSstats calculates log2 fold changes as the ratio of averaged (across replicates) protein intensities between conditions, uses a Student's t-test for p-value calculation and the Benjamini-Hochberg method of FDR estimation to adjust p-values.

### Supplementary Figures

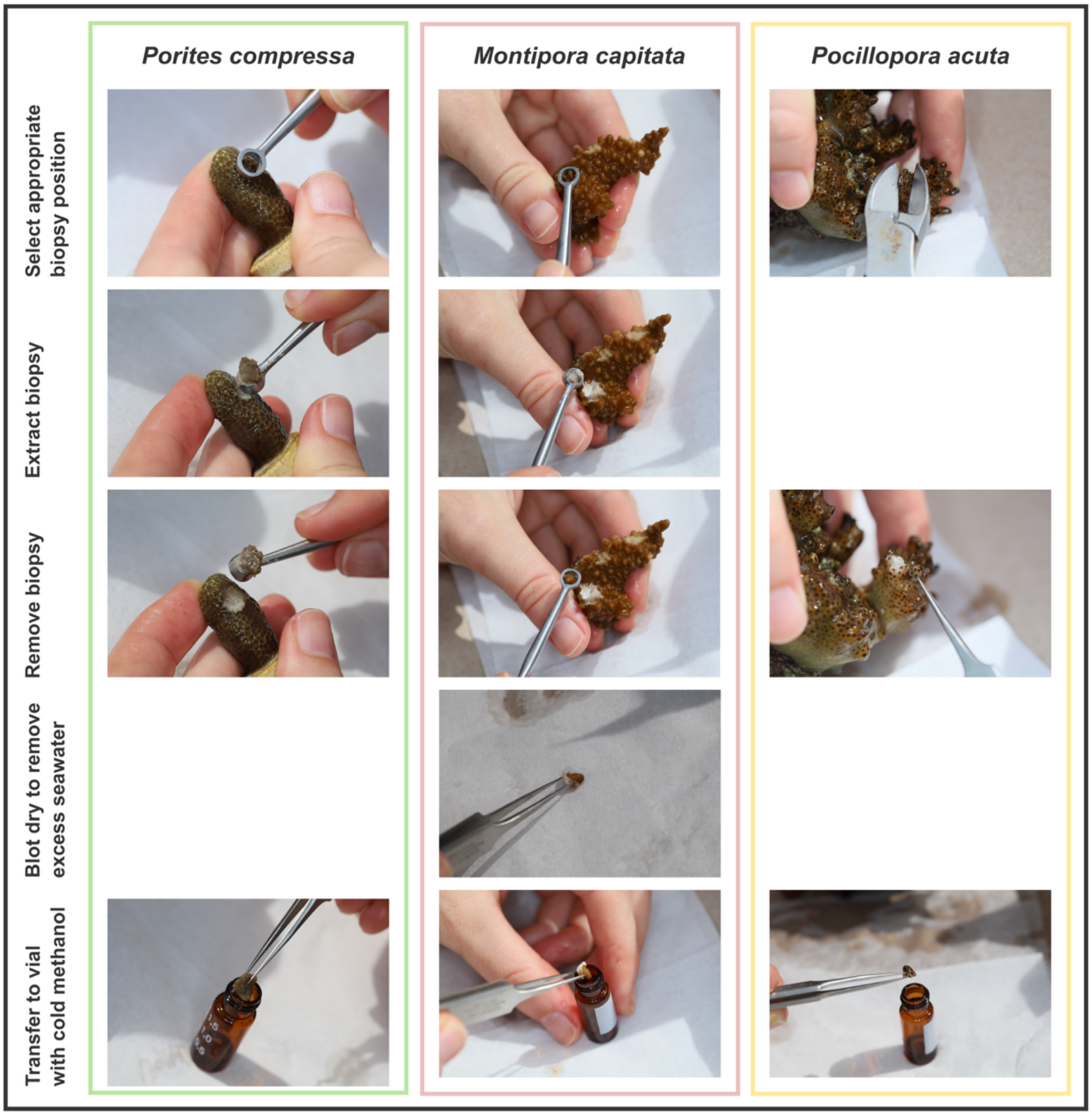


**Figure S1 – Photographs illustrating biopsy collection from three different coral species for metabolomics analysis.** Biopsies are collected from fragments of *Porites compressa* and *Montipora capitata* using a 5mm diameter dermal curette to extract a tiny core of tissue and skeleton by pressing and twisting the curette into the coral fragment and then scooping the biopsy out. Small branches are clipped from a *Pocillopora acuta* colony using bone clippers. The biopsies are briefly blotted dry to remove excess seawater before being transferred into an amber glass vial containing HPLC-grade methanol with 1% formic acid.

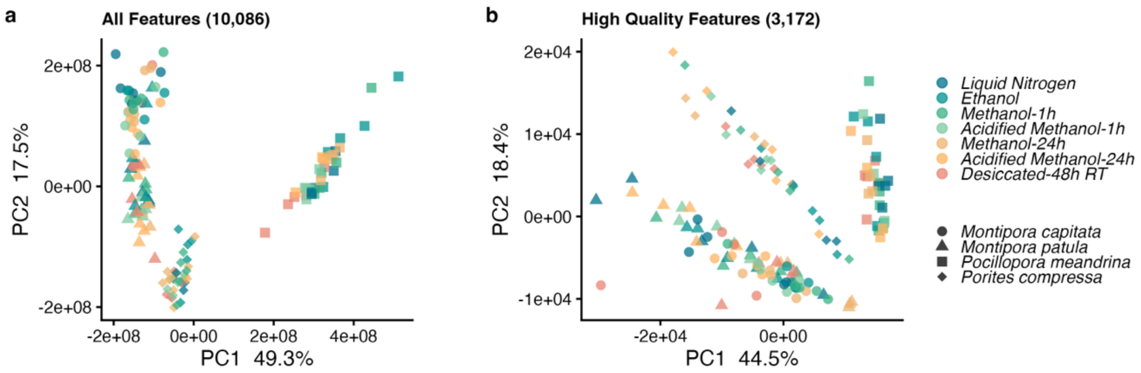


**Figure S2 - Effect of different metabolome quenching and sample handling methods on the holobiont metabolome.** Principle component analysis of **(a)** all detected features and **(b)** high-quality features (thresholds: relative standard deviation for QC samples = 0.4, and Pearson's correlation of serial QC samples = 0.8). Data are normalized using the maximal density fold change (MDFC) method.

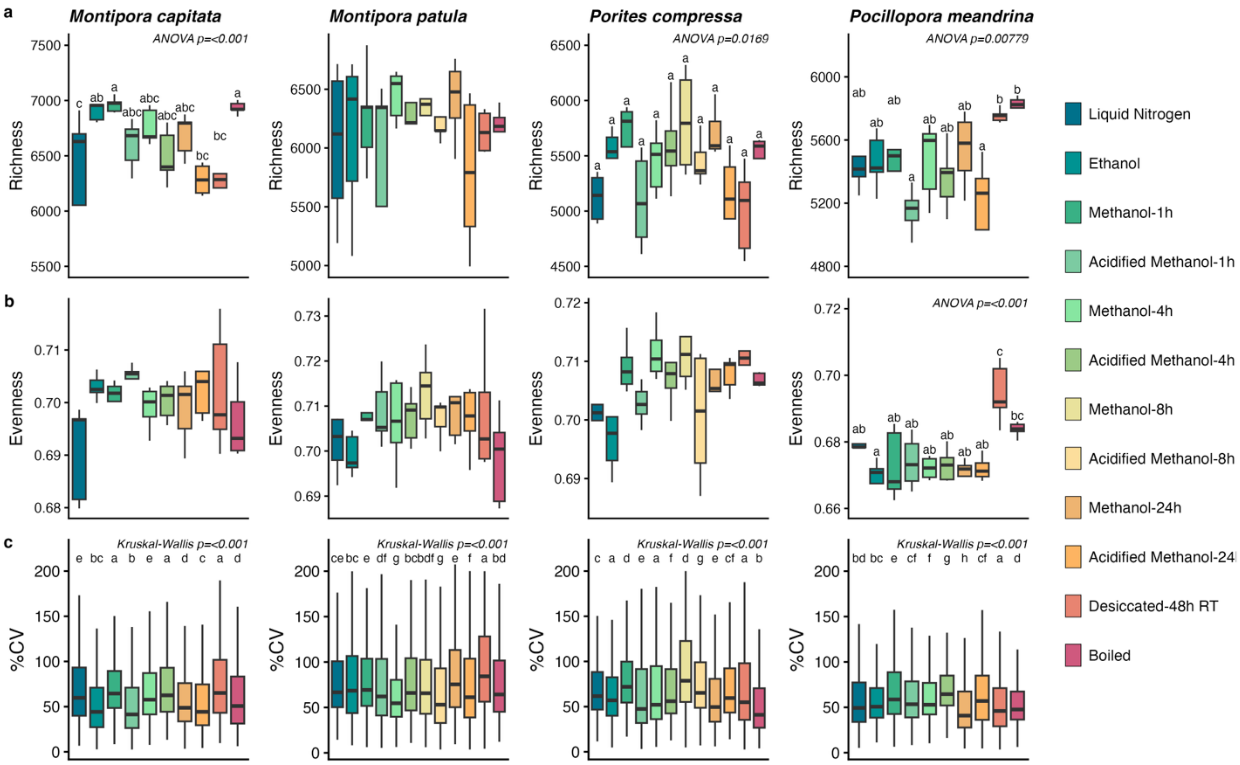


**Figure S3 – Effect of sample fixation and handling methods on metabolome diversity metrics and feature abundance variance. (a)** Metabolite richness, **(b)** evenness, and **(c)** percentage of coefficient of variance (%CV). Analyses were performed on coral holobiont biopsies from *Montipora capitata*, *Montipora patula*, *Porites compressa*, and *Pocillopora meandrina*. Statistical significance was assessed using either one-way ANOVA with Tukey's post hoc test or Kruskal-Wallis with Dunn's post hoc test, depending on data normality.

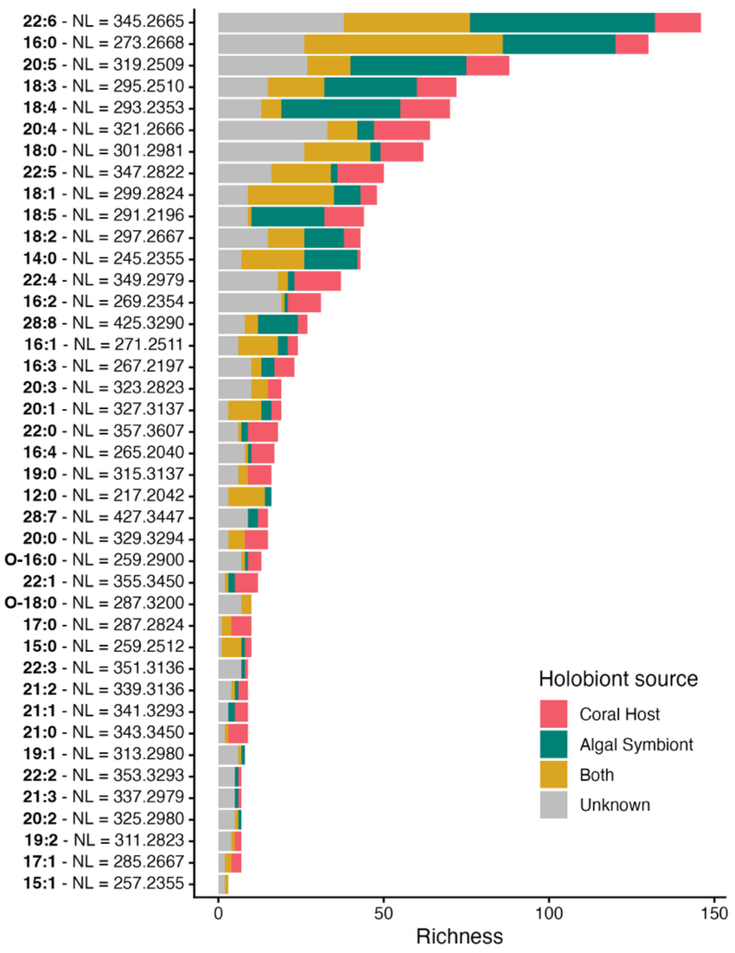


**Figure S4 – Richness of metabolite features with neutral loss (NL) masses corresponding to glycerolipids containing specific fatty acids, and their proportional associations with holobiont compartments**. Metabolite features meeting the requirements of displaying specific NL masses in their $MS^2$ spectra within a mass error of 0.008 were extracted from spectra derived from eight species of coral from Kāneʻohe Bay, Hawaiʻi, and several Symbiodiniaceae genera, both *in hospite* and in culture. In total, these 836 features span an mz range from 298.27-1252.90 and a RT range from 1.2-17.8, and incorporate mono-, di-, and triacylglycerols (MAG, DAG, TAG) as well as glycerolipids containing one ether bond (e.g. mono-alkyl-diacylglycerols – MADAG – designated as O-x). The source of these features within the coral holobiont were determined based on the origin mapping framework presented in the main manuscript.

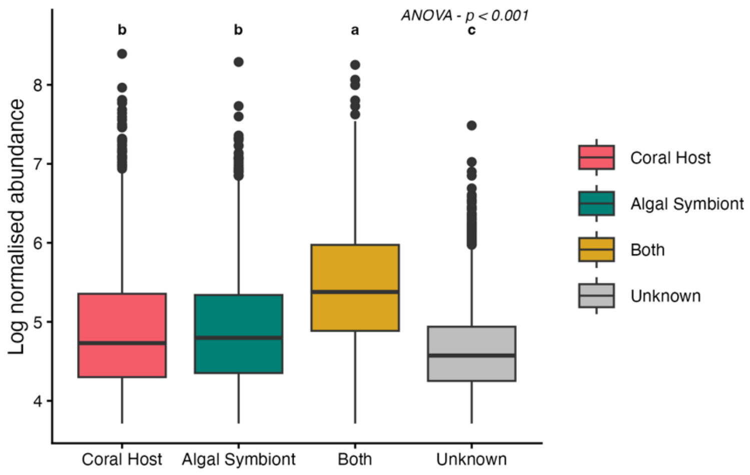


**Figure S5 – Log transformed abundance of all features classified as coral host, algal symbiont, both host and symbiont, or unknown origin** based on mapping feature abundance and presence/absence to representative sample groups of coral host and algal symbiont samples in symbiotic and non-symbiotic or isolated states. Differences between origin classes were assessed by one-way ANOVA, indicating no difference in abundance between coral host and algal symbiont features but that those classified as “Both” had significant increased relative abundance while those classified as “Unknown” had significantly decreased relative abundance, as indicated by the letters.
